# The Marginal Value Theorem in *Caenorhabditis elegans*

**DOI:** 10.64898/2026.08.10.743854

**Authors:** Alid Al-Asmar, Roger Lloret-Cabot, Alfonso Pérez-Escudero

**Affiliations:** Université de Toulouse, Centre National de la Recherche Scientifique, Centre de Recherches sur la Cognition Animale, Centre de Biologie Intégrative, Toulouse 31062, France; Laboratory of Mathematical and Physical Ethology, Research Center of Mathematics for Social Creativity, Research Institute for Electronic Science, Hokkaido University, Sapporo, Hokkaido Prefecture, Japan; Centre d’Estudis Avançats de Blanes (CEAB-CSIC), Cala Sant Francesc 14, 17300 Girona, Spain

**Keywords:** Optimal Foraging *|* Marginal Value Theorem *| C. elegans |* Behavioral Ecology *|* Animal Behavior

## Abstract

The Marginal Value Theorem (MVT) is an important part of Optimal Foraging Theory, predicting the optimal time to leave a food patch. It has been mostly studied in birds, insects and mammals, even though simpler organisms also need to forage efficiently in patchy environments. Here we test whether the nematode *Caenorhabditis elegans* implements the MVT. We recorded individual nematodes exploring patchy environments, across four inter-patch distances and three different food qualities, and found that *C. elegans* behavior matches MVT predictions: When food patches are further away, each food patch is exploited for a longer time. In previous studies animals achieved this by modulating the duration of visits to food patches. Similarly, we found that *C. elegans* also increases visit duration with inter-patch distance, but this only accounts for half of the increase in total exploitation time. The other half of the increase comes from *C. elegans* revisiting food patches multiple times, and the number of these revisits increasing with inter-patch distance. This increase in the number of revisits is not due to behavioral changes in response to distance, but rather to a passive interaction between trajectories and environment geometry. These results show that *C. elegans* can learn the statistics of an environment and use this information in a way consistent with the MVT, but also that part of the fitness-relevant outcomes can emerge passively.

**SIGNIFICANCE:** Despite being key in understanding foraging in patchy resources, the Marginal Value Theorem (MVT) has been tested almost exclusively in relatively complex animals. We extensively tested the MVT in a simple, non-visual organism, showing that *Caenorhabditis elegans* increases patch exploitation time when inter-patch distance increases. This effect is partially driven by the same behavioral adaptation found in complex animals, but also by an increase in the number of patch revisits. This second driver, which had not been reported before and is probably key for non-visual organisms, requires no behavioral adaptation and produces around half of the fitness-relevant outcome. Our results highlight the need for adapting Optimal Foraging Theory to a wide range of taxa spanning from microbes to small invertebrates.

## INTRODUCTION

Optimal Foraging Theory provides tools to predict and assess how animals find and consume food (Ritvo, 2022; D. W. Stephens and Krebs, 1986). While organisms should not be expected to be perfectly optimal and often deviate from the optimum in systematic ways (Chen et al., 2006; De Petrillo and Rosati, 2019; Lea and Ryan, 2015; Nonacs, 2001; Parker and Smith, 1990; Pérez-Escudero et al., 2009; Shettleworth, 2010), Optimal Foraging Theory provides a useful reference and has been successful in explaining many features of animal behavior (Ritvo, 2022; D. W. Stephens and Krebs, 1986). A key question is the optimal way of exploiting heterogeneous environments, where resources are patchily distributed. Historically, this question has been addressed using the Marginal Value Theorem (MVT), which prescribes the optimal time to leave a patch of food (Charnov, 1976). One of the main predictions of the MVT is that the optimal time to leave a food patch depends on the quality and distance of other food patches, requiring the individual to form an overall estimate of the environment, rather than to just focus on the current food source. This prediction has been tested in many insects and vertebrates (Cassini et al., 1990, 1993; Giraldeau and Kramer, 1982; Kacelnik and Todd, 1992; Marshall et al., 2013; Nonacs, 2001; O’Bryan et al., 2020; Pleasants, 1989; Pyke, 1978; Tenhumberg et al., 2001; Wajnberg et al., 2000; Watanabe et al., 2014; Zimmerman, 1981), including humans (Bella-Fernández et al., 2022; Louâpre et al., 2010; Pacheco-Cobos et al., 2019; Schlender et al., 2024). However, all these studies focus on animals with comparatively complex sensory and nervous systems, such as insects, birds or mammals.

Simpler organisms, and in particular those without vision, face the same foraging challenges and must address them with stronger cognitive and sensory constraints. Certain aspects of foraging have been extensively studied in microorganisms and small invertebrates, including search (Hills et al., 2004; Salvador et al., 2014; Zjacic and Scholz, 2022), chemotaxis (Berg, 2004; Iino and Yoshida, 2009; Suwazono et al., 2025), food choice (Dussutour et al., 2019; Fukasawa and Ishii, 2023; Katzen et al., 2023; Madirolas et al., 2023; Shtonda and Avery, 2006), patch-leaving, (Haley et al., 2025; Latty and Beekman, 2009; Milward et al., 2011; Scheer and Bargmann, 2023), dispersal (Donahue et al., 2003) sensory adaptation (Colbert and Bargmann, 1995; Dekkers et al., 2021) and learning (Boussard et al., 2019; Cho et al., 2016; Dussutour, 2021; Saeki et al., 2001). However, the main prediction of the MVT, which links patch residence time to the distance and quality of other food patches, has rarely been tested in these systems (Hohberg and Traunspurger, 2009). Besides their ecological importance, small, short-lived organisms offer an opportunity to test Optimal Foraging models rigorously, because they can be understood with great mechanistic detail and studied with large numbers of individuals, long distances relative to body size and long experimental times relative to lifespan (Al-Asmar and Pérez-Escudero, 2026; Anderson and Perona, 2014).

To take advantage of these opportunities and improve our understanding of Optimal Foraging in simple organisms, we turned to the nematode *Caenorhabditis elegans*, a 1 mm-long worm with 959 cells, including 302 neurons, which feeds on the bacteria that grow on decaying organic matter (Al-Asmar and Pérez-Escudero, 2026). Owing to its small size, fast life cycle, and ease of cultivation, *C. elegans* is a widely used model organism (Dietrich et al., 2014; Nigon and Félix, 2018). Its foraging-related behaviors have been extensively studied, including locomotion (Ahamed et al., 2021; Costa et al., 2024; Gomez-Marin et al., 2016; Schwarz et al., 2015; G. J. Stephens et al., 2008, 2011; Yemini et al., 2013), search (Hills et al., 2004; Salvador et al., 2014), chemotaxis (Iino and Yoshida, 2009; Pierce-Shimomura et al., 1999; Suwazono et al., 2025; Ward, 1973), food choice (Madirolas et al., 2023; Shtonda and Avery, 2006), feeding (Arous et al., 2009; Ding et al., 2020; Lee et al., 2017), patch handling (Ding et al., 2020; Haley et al., 2025; Iwanir et al., 2016; Milward et al., 2011), collective patterns (Ding et al., 2019; Perez et al., 2025), associative learning (Gourgou et al., 2021; Saeki et al., 2001; Tasnim et al., 2025), and behavioral states (Arous et al., 2009; Scheer and Bargmann, 2023), although most studies have been performed in highly stereotyped laboratory conditions aimed to dissect mechanistic details of *C. elegans* behavior, rather than to understand its function in an ecological context (Al-Asmar and Pérez-Escudero, 2026; Frézal and Félix, 2015; Petersen et al., 2015). Several studies have looked into *C. elegans* foraging in patchy environments, finding that worms facing a novel environment tend to explore several food patches before settling to exploit one of them (Haley et al., 2025; Pradhan et al., 2019), prefer high-quality food patches regardless of their species composition (Madirolas et al., 2023), and leave food patches once they are depleted (Milward et al., 2011). While these results have shown that the worms can modulate their response to food based on both local stimuli and past experience, it has never been tested whether this modulation is enough for worms to adapt to large-scale environmental features such as inter-patch distance, especially on a long time scale.

Here, we test whether the MVT applies to *C. elegans* by studying its foraging behavior in a patchy environment. We recorded more than a thousand worms foraging individually in arenas with different inter-patch distances and food qualities. This approach allows us to quantify how patch residence times depend on both food distribution and quality, and to assess whether the nematode’s behavior follows the predictions of Optimal Foraging Theory.

## RESULTS

### An experimental model for foraging in patches in *C. elegans*

We developed a high-throughput pipeline to record worms while they are foraging on patches of bacteria. In each experimental replicate, a single worm was placed in a 55 mm diameter arena, where food patches had been placed the previous day following a triangular grid pattern (Figure 1A). Food patches consisted of small (0.75 µL) drops of *E. coli* OP50, so that each food patch could be depleted in a relatively short time. We used a pipetting robot to increase the accuracy and repeatability of the food patches, which had a radius of 1.4 mm *±* 0.2 and a standard deviation of 2% in inter-patch distance (Figure S1H-I). We used oblique illumination to determine the exact position and geometry of each food patch, locating patch edges with an accuracy of 1 pixel (0.03 mm) (Figure S2). Each non-control plate had one of three food qualities (OD = 0.2, OD = 0.5 and OD = 1.25, measured as Optical Density at 600 nm), with control plates containing drops of buffer instead of bacteria (hereafter OD = 0). Our experimental plates were optimized to prevent bacterial growth, ensuring that worms were exposed to the target bacterial qualities (Madirolas et al., 2023). We recorded each worm for 8 hours, and analyzed the video to obtain the worm’s silhouette and its center of mass in every frame.

**Figure 1:**
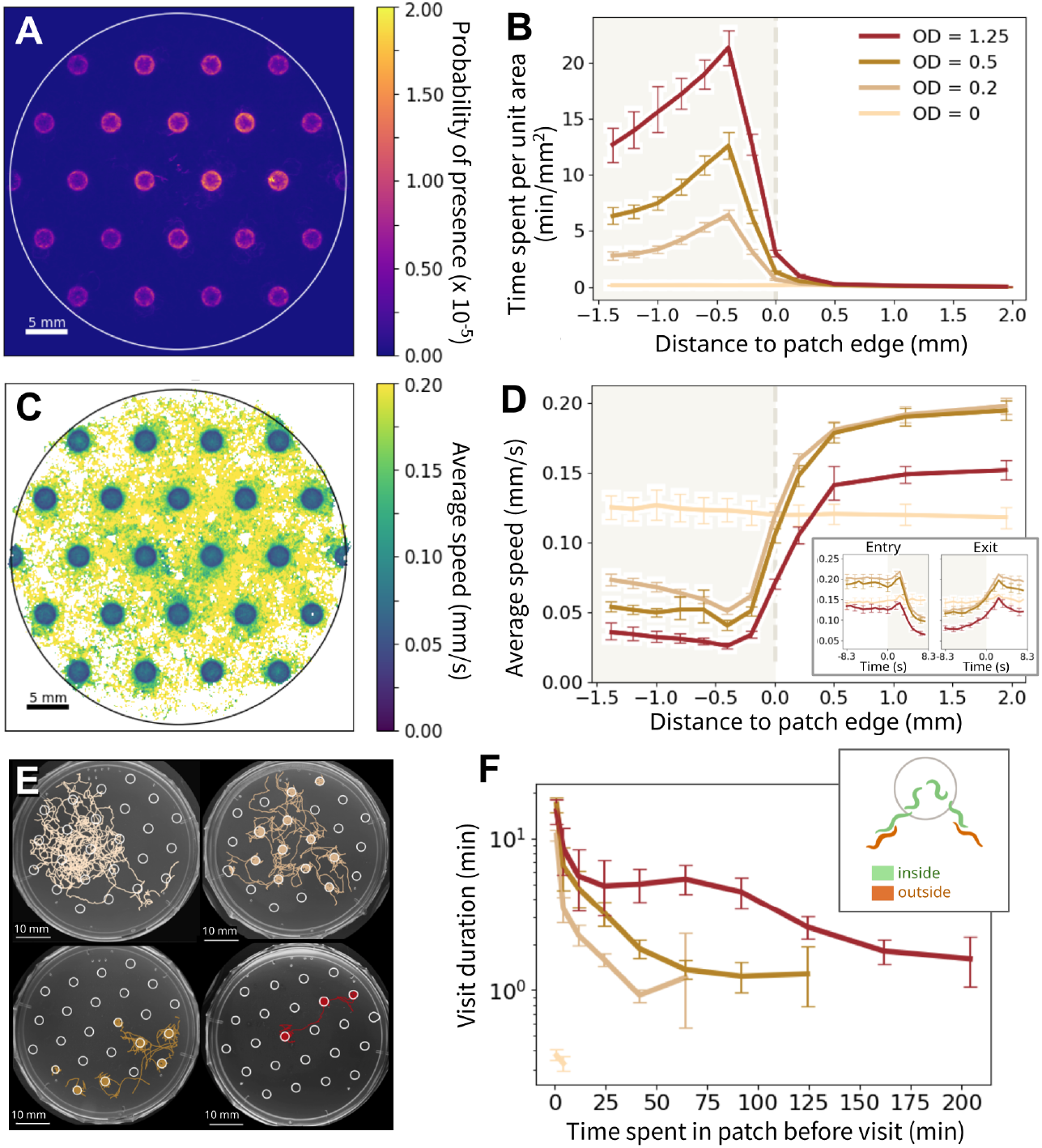
Overview of the worms’ response to our experimental setup. **A:** Probability of presence of the worm in all pixels of the arena, for Optical Density OD = 0.2. Coordinates have been transformed for patch locations and sizes to exactly match between plates (see Methods). In these transformed coordinates, the region of interest (ROI) for the tracking is slightly different for each plate. White line shows an approximate average ROI. **B:** Average time spent per unit area as a function of distance to the edge of the food patch (negative distances inside the food patch). Each line corresponds to a food quality (OD of the bacteria). **C, D:** Same as A, B but for the average speed. **Inset:** Speed of the worm as a function of time around patch entry (left) or patch exit (right) (only keeping entry/exits when the worm remained inside the food patch for at least 8.3 seconds before the entry/exit and for at least 8.3 seconds after it). **E:** Examples of worm trajectories. The color of each trajectory indicates food quality (OD, see other panels for legend). White: bacterial food patches. **F:** Average duration of the visits to a patch as a function of the time spent in that food patch before the visit (i.e. the first visit to a food patch is at time 0, the second visit is counted at the time duration of the first visit, etc.). Each curve corresponds to a food quality. **Inset:** Definition of visits: the worm is considered to be inside a food patch whenever any of its pixels overlaps with the patch. **All boxes:** Errorbars are 95% bootstrap confidence intervals (1000 resamples, see Methods).

Worms responded to the food patches in a way consistent with previous studies (Iwanir et al., 2016; Scheer and Bargmann, 2023): They spent much more time inside the food patches than outside them, showing a preference for food patch edges (Figure 1A, B), and they moved more slowly when inside (Figure 1C, D). Both of these effects were stronger for higher food quality (Figure 1B, D).

We defined a visit event as starting when any part of the worm touches a food patch, and ending once the worm body has completely left (Figure 1F, inset). Worms visited multiple, but rarely all patches within the experimental time (Figure 1E and Figure S3). They often visited the same food patch multiple times, and we used this fact to study the effect of food depletion: Most theoretical models assume that food patches provide diminishing returns, giving out less and less food per unit of time as they get depleted (Charnov, 1976; D. W. Stephens and Krebs, 1986). Given that worms respond to lower food quality by reducing the time spent on the patch (Figure 1B), if our food patches indeed provide diminishing returns, revisits to a food patch should be shorter, on average, the longer the worm has spent previously on that patch. This is what we found, with visit duration decreasing from around 20 min (fresh food patch) to just a couple of minutes (Figure 1F). This shortest visit duration is reached earlier the lower the initial quality. These results are consistent with a previous study that found that leaving rate increases progressively on large food patches populated by a group of worms (Milward et al., 2011).

### *C. elegans* spends more time in each patch as inter-patch distance increases

We next tested the main prediction of the Marginal Value Theorem (MVT), which states that an individual should leave a patch once their feeding rate in that patch has reached the average expected feeding rate in their environment (Charnov, 1976). A way to test this theorem is to change the distance between food patches, because higher inter-patch distances lead to higher transit times, and thus to a lower average feeding rate. Therefore, according to the MVT, when the distance between the food patches increases, the average residency time in each patch should increase (Figure 2A). This prediction is interesting because implementing it requires individuals to integrate local and global information (current food, but also inter-patch distance). It is especially interesting in organisms lacking vision such as *C. elegans*, as they cannot trivially estimate global information such as distance to a non-visited patch.

**Figure 2:**
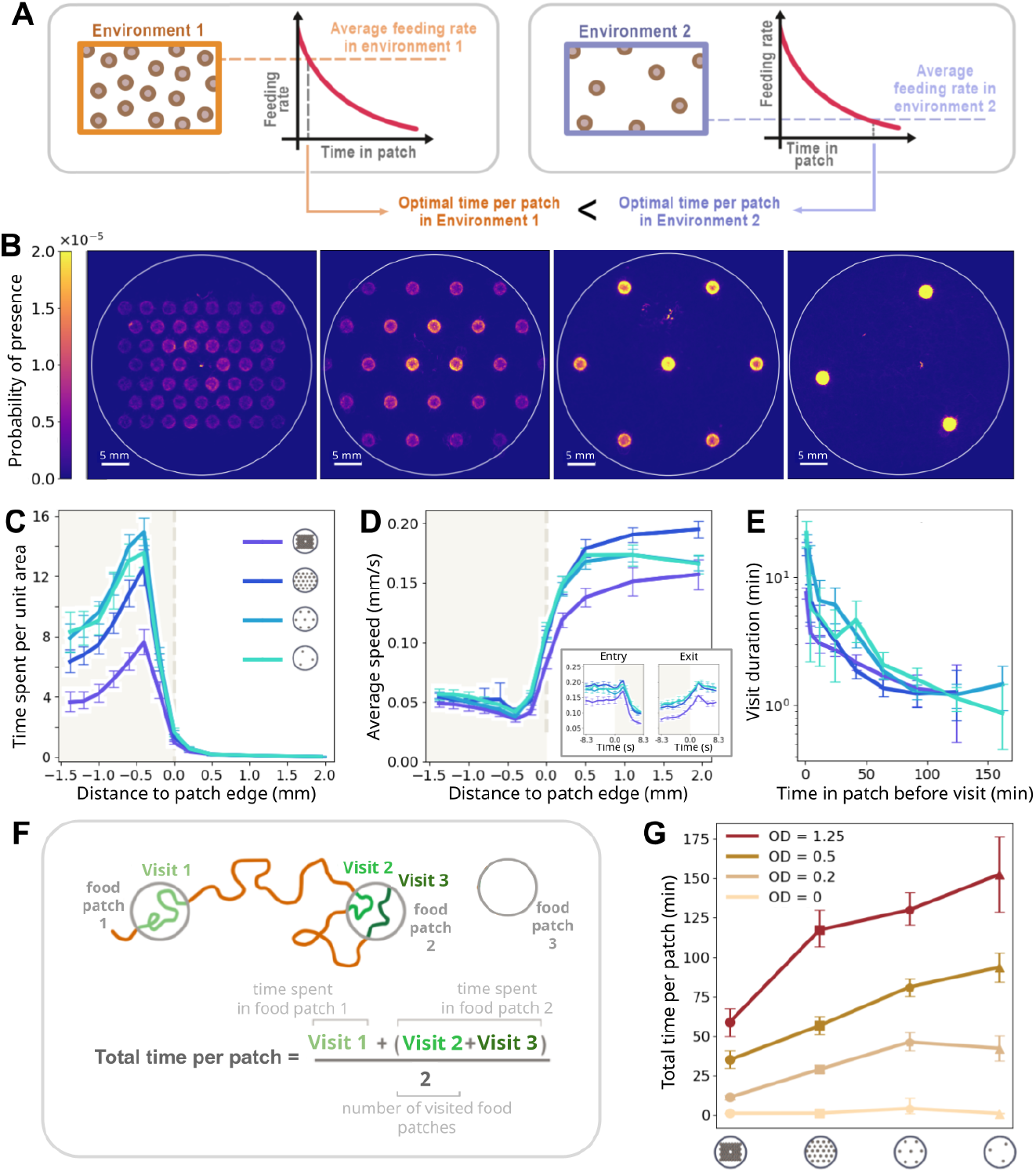
The time spent in each patch increases with inter-patch distance, as predicted by the MVT. **A:** Main result of the MVT: Environment 1 (left) has closer food patches than Environmen 2 (right), so it has a higher average feeding rate (dashed lines). The plots show the feeding rate experienced by an animal on a food patch, which decreases over time due to diminishing returns, and is identical in both environments. The MVT dictates that the animal should leave the food patch when the feeding rate matches the environmental average, predicting a longer optimal time in patches in Environment 2 than in Environment 1. **B-E:** Same as Figure 1A, B, D, F, for all inter-patch distances and for quality OD = 0.5. See Figures S4, S5, S6, S7, S8 for full datasets and alternative normalizations. **F:** Top: Sketch of trajectory, indicating transits and visits to food patches. Bottom: Equation for the total time spent per patch. **G:** Total time per patch as a function of inter-patch distance. Each line corresponds to one food quality. **All boxes:** Error bars are the 95% bootstrap confidence intervals (1000 resamples, see Methods).

To test this prediction in *C. elegans*, we replicated the setup shown in Figure 1, but with four different inter-patch distances, ranging from 4.5 to 28 mm (Figure 2B). First, we checked that the results presented in Figure 1 hold qualitatively for all distances: In all cases worms spend more time inside the food patches and particularly near their edge (Figure 2C), slow down inside the food patches (Figure 2D) and decrease the visit duration in subsequent revisits, in a fashion compatible with diminishing returns (Figure 2E) (see full results in Figures S5, S4, S6, S8).

The quantity of interest in the MVT is the total time spent in each food patch, as it determines the extent to which a food patch has been depleted. We measured the total amount of time that the worms spent in each visited food patch during our experiment (Figure 2F), and found that it increased as distance increased, as predicted by the MVT (Figure 2G). However, it was not immediately clear what behavioral changes drove this increase in total time per patch: The speed inside the food patches did not decrease with inter-patch distance (Figure 2D) and visit duration dynamics were similar across distances (Figure 2E).

### The increase in total time per patch is only partially driven by behavioral changes inside food patches

In our experiments, *C. elegans* visits each food patch multiple times. This means that the total time spent in each patch depends on: (1) the number of visits, and (2) the duration of each individual visit (Figure 3A). In all conditions, the number of visits per patch increased with distance (Figure 3B, S9). By contrast, the average visit duration increased with inter-patch distance only in one of the tested food qualities (OD = 0.2). In the two other conditions (OD = 0.5 and OD = 1.25), no increase was observed (Figure 3C, S9).

**Figure 3:**
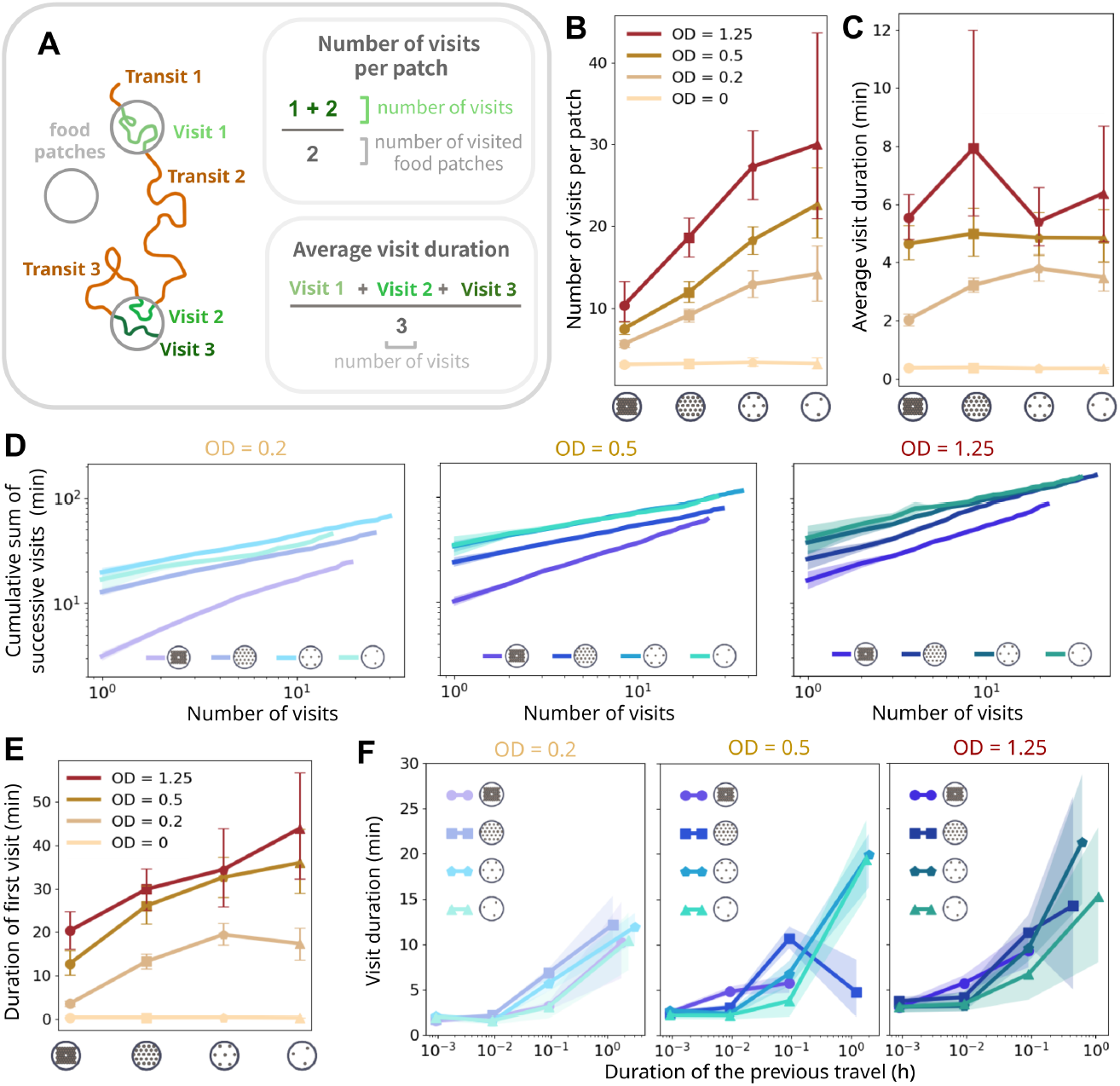
Behavioral adaptation to inter-patch distance. **A:** Left: Sketch of trajectory, indicating transits and visits to food patches. Right: Equations for the number of visits per patch and the average visit duration. **B:** Number of visits per patch, as a function of inter-patch distance. Each line corresponds to one food quality. **C:** Same as B, but for average visit duration. **D:** Cumulative time spent in a food patch up to (and including) the *n*-th visit, as a function of *n*. These times are computed by averaging the times of all first, second, third, etc. visits, and then computing the cumulative sum of these averages. Each box corresponds to one food quality, and each line corresponds to one inter-patch distance. **E:** Same as B, but for duration of the first visit to each food patch. **F:** Visit duration as a function of the duration of the preceding transit (points with less than N = 10 visits are not shown). **All boxes:** Errorbars and shaded areas represent 95% bootstrap confidence intervals (1000 resample, see Methods).

However, average visit duration is obscured by the effect of diminishing returns: Visits become shorter as a food patch gets revisited multiple times (Figure 1F), and patches that receive more visits accumulate more of the shorter late visits, which drives their average visit duration downwards. In order to control for this effect, we looked at how the total time spent in a patch increases as successive visits accumulate. We found that this cumulative time grows following sub-linear power laws, and shows clear differences across inter-patch distances (Figure 3D). The clearest difference can be seen in the vertical shift of the curves, which represents the first visit to each food patch. This first visit is by definition not affected by patch depletion, and its duration increases both with food quality and inter-patch distance (Figure 3E). These results show that the worms do alter their behavioral response to food as a function of the distance between patches, in a way consistent with the MVT (longer visit durations in poorer environments). Since worms cannot trivially estimate inter-patch distances, we hypothesized that they might be responding to the duration of the previous transit to modulate visit duration. We found that the longer a transit is, the longer worms remain a food patch when they encounter one (Figure 3F). Transit duration could thus be used as a proxy of inter-patch distance and, more generally, of environmental quality.

### The increase in number of visits is not driven by behavioral adaptation to interpatch distance

In higher inter-patch distances, each patch gets visited more times on average (Figure 3B). This increase in number of visits may be simply due to the fact that higher inter-patch distances mean fewer patches in the arena, so the worm has more time to revisit each of them, but it could also be a consequence of behavioral changes in response to different inter-patch distances. In particular, worms could be altering their speed and trajectories after exiting a food patch, which could change their probability of coming back to it. In order to test for this effect, we looked at the probability of worms reaching a certain distance from the patch edge after leaving (Figure 4A), and at the average time it took worms to reach that distance (Figure 4B). We found no clear difference between distances in almost all conditions, suggesting that changing interpatch distance does not change behavior outside the food patches in a way that affects revisit probabilities.

**Figure 4:**
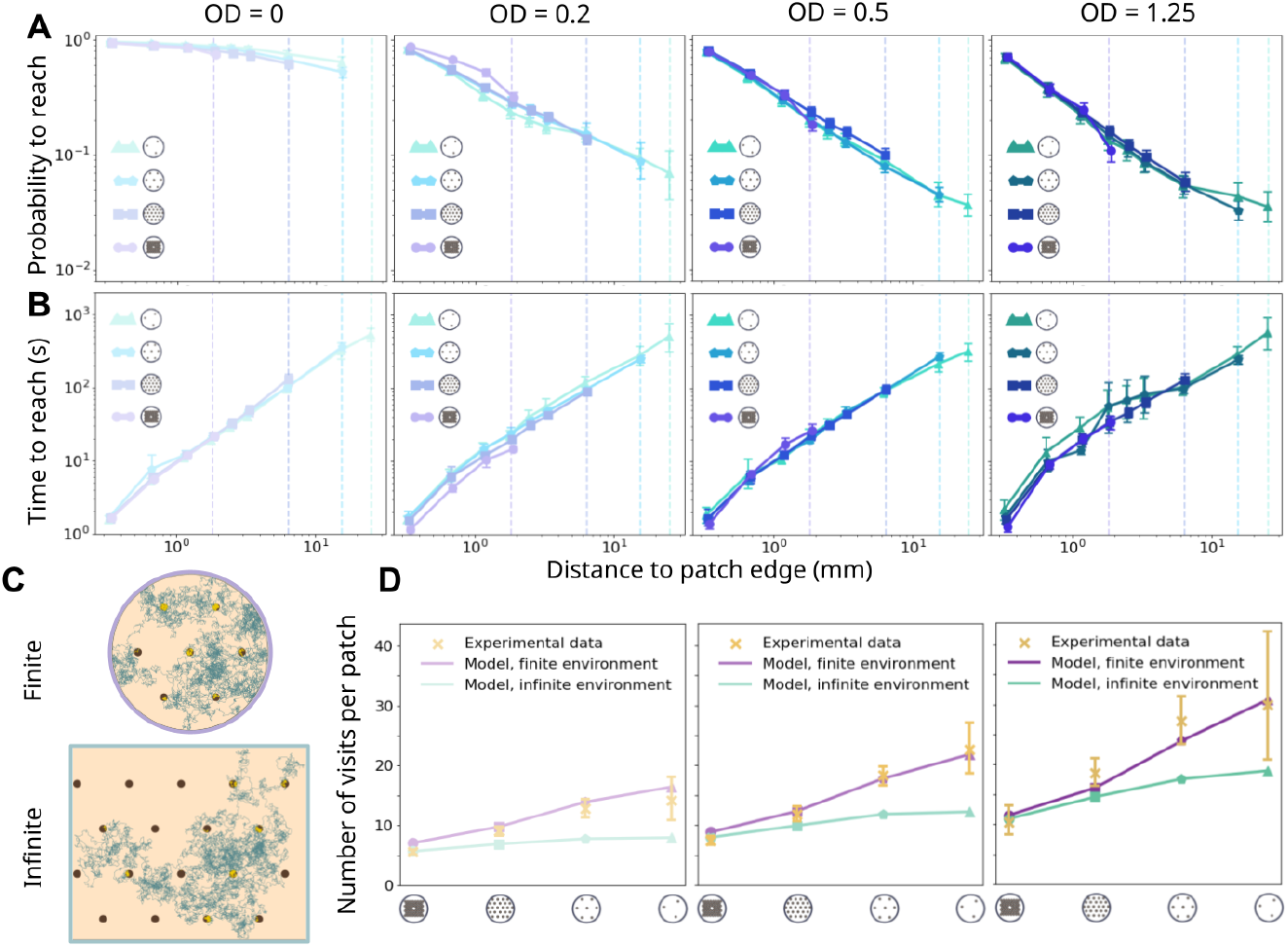
The increase in number of visits is not driven by behavioral adaptation to inter-patch distance. **A:** Probability that the worm reaches a given distance to the patch edge after exiting a patch. Each color corresponds to an inter-patch distance, and data are shown only for distances below the distance to the nearest patch (vertical lines). Each column corresponds to a food quality. **B:** Same as A, but for the time taken by the worm to reach a given distance to the patch edge after exiting a patch. **C:** Simulation of random walk trajectories with 18 mm interpatch distance, for finite and infinite environments. **D:** Average number of visits to each food patch, as a function of inter-patch distance, for our experiments (yellow dots) and for simulations (1000 agents, and 8 hours) in finite (purple) and infinite (green) environments. **All boxes:** Error bars are the 95% bootstrap confidence intervals (1000 resamples, see Methods).

To study how the increase in number of visits happens without behavioral adaptation to interpatch distance, we simulated agents that do not change their behavior as a function of inter-patch distance, in environments with the same patch arrangements as our experiments. We used a random walk model, in which agents reorient themselves randomly at some temporal frequency, and progress at two different speeds depending on whether they are inside, or outside food patches (Figure 4C). We fitted the three model parameters to best reproduce number, average duration, and sum of visits made to each visited food patch in our experiments (Supplementary Table S3). The fitting was made once for each food quality, without changing the parameters across inter-patch distances.

This simple model reproduces the increase in number of visits as a function of inter-patch distance (Figure 4D, purple curves), showing that behavioral adaptation to inter-patch distance is not necessary to reproduce the phenomenon. This means that it could instead be the result of an interaction between the worm’s behavioral programme (which does not change with distance), and the geometry of the environment.

This increase in number of visits emerges from the finite nature of our experiments, both in space and in time. Finite space means that higher inter-patch distance leads to fewer food patches, each of them accumulating more visits. But even in an infinite space we can observe an increase of number of visits with inter-patch visits, provided that time is finite: Smaller interpatch distances mean that more food patches are found between successive revisits to a given food patch. Since each visit to another food patch takes extra time, it increases the probability that the experiment finishes before the original food patch is revisited. To illustrate this effect we simulated the same agents in infinite environments, showing that the increase in number of visits with inter-patch distance is diminished but conserved (Figure 4D, green curves).

### Behavioral adaptation and passive effects contribute similarly to increasing total time per patch with inter-patch distance

Total time per patch increases as inter-patch distance increases, and this is driven by both the duration of visits, and their number (Figure 3). While the visit duration increase can be attributed to a behavioral response, the number of visits seems to increase regardless of whether worms change their out-of-patch behavior (Figure 4). Because of this lack of behavioral change, we refer to this second factor as passive. We have built a model to disentangle the relative contribution of in-patch effects and out-of-patch passive effects.

To capture out-of-patch effects, we first characterized the transition matrix across food patches, computing the probability that a worm moves from one food patch to another (Figure 5A). This transition matrix includes experimental complexities, such as how worms are more likely to return to the patch they have just exited, and how peripheral patches differ from central ones. Then, we characterized the duration of these transitions, again building a matrix that distinguishes transitions for every pair of food patches (Figure 5B). Each element of this matrix actually represents a whole distribution of transit durations (see Methods for details). Similarly, to describe the beginning of the experiment, we built a first-transit array, containing the probability to reach each patch first and the distribution of durations it took to reach them.

**Figure 5:**
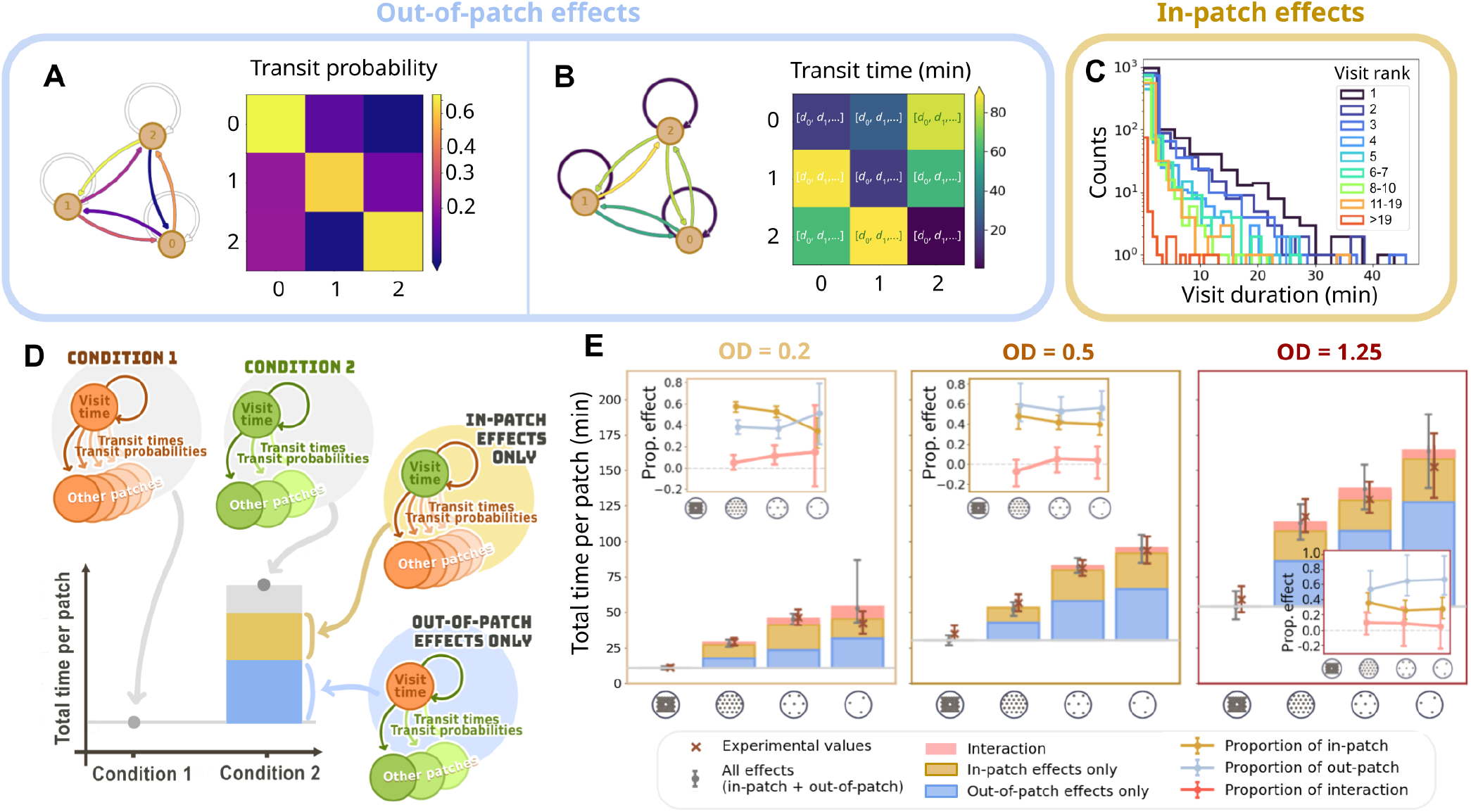
Relative contribution of in-patch and out-of-patch effects. **A:** Transition matrix showing the probability of a transit between any two patches, for the three-patch condition with quality OD = 0.2. Arrow colors show transition probability, using the same scale as the heatmap. See all matrices in Supplementary Figure S12 **B:** Same as A, but for the average transit time between every two patches. Lists in each element of the matrix represent the whole list of used to draw transit times in the model. **C:** Histogram of visit durations for the three-patch condition with quality OD = 0.2. Colors represent visit rank (1st visit, 2nd visit, etc.). **D:** Schematic of our model and the plots in panel E. Condition 1 is the baseline condition (orange). In the full model, condition 2 is simulated using its visit durations and transits (green), and its result is represented by the gray dot. To simulate out-of-patch effects only, the model uses visit durations from condition 1 and transitions from condition 2 (cartoon with blue background), and the result is shown by the blue bar. To simulate in-patch effects only, the model uses visit durations from the condition 2 and transits from the condition 1 (cartoon with yellow background), and the result is shown by the yellow bar, which is stacked on top of the blue bar. The difference between the full model and the addition of the two separate effects (red bar) represents the interaction between out-of-patch and in-patch effects. **E:** Total time per patch as a function of inter-patch distance, and for each food quality. Orange crosses: Experimental values. Gray dots: Full model. Bars: Contribution of each component of the model, as described in panel E. Insets: Relative contribution of each type of effect, computed as follows: Let *F*, *O*, *I* be the total time per patch computed using the Full model, Out-of-patch effects only, and In-patch effects only, respectively. The blue dots show *O/F*, the yellow dots show *I/F*, and the red dots show (*F − O − I*)*/F*. Error bars are the 95% bootstrap confidence intervals (1000 resamples, see Methods).

To capture in-patch effects, we extracted visit durations, assuming that they depend on the number of visits already made to the food patch, and that their distribution is equal for all food patches in a given condition (Figure 5C).

We then modeled the behavior of a worm as follows: The worm enters a patch chosen with a probability given by the first-transit matrix and at a time extracted from the first-transit duration distribution. Then, it spends a time in the patch extracted from the corresponding distribution of visit durations (in this case, corresponding to a first visit). It then transits to another patch (which can be the same as the original one), with probabilities taken from the transition matrix, and spending a transit time taken from the corresponding distribution in the transit-time matrix. The simulation continues in this way until it reaches the experimental time of 8 hours (see Methods for a more detailed description).

We ran this model for all conditions, obtaining good agreement with the experimental data (Figure 5D, E, gray dots). Then, we used the model to disentangle the contribution of in-patch and out-of-patch effects: Using the condition with smallest inter-patch distance as a reference, we simulated the effect of applying the out-of-patch effects (i.e. transition matrices) from the other conditions, while keeping the in-patch effects (i.e. visit durations) from the reference condition (Figure 5D, blue). In this way, this simulation mimics a worm that does not change visit duration with inter-patch distance, retaining only out-of-patch passive effects. This produced an increase in total time per patch roughly half of the one observed experimentally (Figure 5E, blue bars). Conversely, we performed a simulation where only the in-patch effects are present, by keeping the out-of-patch effects of the reference condition and changing only visit duration with interpatch distance (Figure 5D, E, yellow bars). The combination of out-of-patch and in-patch effects almost amounts to the results of the full simulation, with the difference corresponding to the interaction between the two effects, which is absent when running simulations with one effect only (Figure 5E, red bars).

We then computed the proportion of the increase in total time per patch due to each separate effect (Figure 5E, insets), finding that this interaction is not statistically significant in most cases (i.e. the two effects are additive), and that the relative contribution of out-of-patch effects increased with food quality.

## DISCUSSION

We have tested the Marginal Value Theorem (MVT) on *C. elegans*, finding that worms can adapt their behavior to large-scale environmental features, such as inter-patch distance. The relevant outcome of this adaptation—the increase in total time spent in each patch when inter-patch distance increases—is a combination of two effects that have similar weight: A behavioral adaptation to increase visit duration, which requires the worms to perceive and respond to the characteristics of the environment, and passive effects arising from the interaction between movement trajectories and environmental geometry.

The behavioral adaptation leading to longer visits is compatible with previous work that showed that worms that had fed on high food qualities in the past move faster (Sawin et al., 2000), or are more likely to not react (Haley et al., 2025) when encountering lower quality food. By extension, in our setup where satiety is modulated by inter-patch distance rather than food quality, one could hypothesize that when food patches are closer and worms experience food more frequently, they react less strongly to food patches, moving overall faster. However, on average, we only observed this effect for the lowest food quality, while for the two highest food qualities the speed inside the food patches was constant, or even lower for the closest inter-patch distance at OD = 0.5 (Supplementary Figure S7), which suggest a non-trivial interaction between food quality and distance. While previous results were obtained by conditioning worms for long periods of time before the experiment (Haley et al., 2025; Sawin et al., 2000), our study shows that worms can tune their response to food patches in response to a large-scale environmental feature, which only affects indirectly the amount of food they encounter at each point in time. This suggests that *C. elegans* learning is fast enough to adapt to new environments of this size, while being slow enough to average out the fluctuations experienced when exploring different food patches.

Similarly to results found in birds (Houston, 2009; Nonacs, 2001), we also found that visit durations could be in part predicted by the duration of the preceding transit time (3), suggesting worms integrate past feeding experience to modulate their visit durations. One important difference between worms and birds however, is that worms can probably not control directly their leaving of a patch, rather leaving stochastically, at a rate which also depends on the rate at which they encounter the patch edge. While the consequences of stochastic leaving of food patches has been studied from the point of view of optimality (Adler and Kotar, 1999), there has been little work on how worms actually modulate this leaving rate (Scheer and Bargmann, 2023), a behavior which is hard to disentangle from movement speed, as the faster a worm moves, the more chance it has to encounter the patch edge.

The second effect contributing to increasing the total time per patch when increasing interpatch distance is an increase in the number of visits to each patch. While this effect contributes around half of the total outcome, it had been largely neglected in previous works on the MVT: Most experimental works either make revisits impossible or irrelevant by design (Cassini et al., 1990, 1993; Giraldeau and Kramer, 1982; Kacelnik and Todd, 1992; O’Bryan et al., 2020; Wajnberg et al., 2000), study situations where food patches are not clearly defined (Pacheco-Cobos et al., 2019; Watanabe et al., 2014), report negligible revisit rates (Pleasants, 1989; Schlender et al., 2024), or do not mention revisits (Marshall et al., 2013; Pyke, 1978). Some studies found significant revisit rates, but did not study the change in revisit rate with inter-patch distance (Hong and Wolfe, 2023; Tenhumberg et al., 2001; Zimmerman, 1981). In contrast, our results show that revisits play a key role for *C. elegans*. They probably also play an important role in many other organisms, especially those with limited spatial awareness whose search patterns inevitably lead to revisiting the same locations.

The consequences of revisiting previously explored space have been extensively studied in random search theory. This literature has found efficient search strategies that balance intensive sampling of recently profitable regions with extensive exploration of unvisited space. This balance can be achieved through several mechanisms, including scale-free step-length distributions such as in Lévy walks (Bartumeus et al., 2002; Bartumeus et al., 2005, 2014; Viswanathan et al., 1999; Viswanathan et al., 2011), correlated or composite random walks (Benhamou, 2007, 2014; Codling et al., 2008), intermittent search strategies that alternate between local detection and relocation phases (Bénichou et al., 2011; Nolting et al., 2015), and area-restricted search, in which recent resource encounters or expectations increase local search effort (Bartumeus and Levin, 2008; Benhamou, 1992; Dorfman et al., 2022). These strategies differ in the degree to which they reduce oversampling, promote returns to resource-rich regions, or accelerate displacement to new areas.

While this body of work has provided substantial insight into how movement strategies influence encounter rates (Bartumeus et al., 2002; Bartumeus and Levin, 2008; Bartumeus et al., 2005, 2014; Benhamou, 1992, 2007, 2014; Bénichou et al., 2011; Codling et al., 2008; Dorfman et al., 2022; Nolting et al., 2015; Viswanathan et al., 1999; Viswanathan et al., 2011), our results highlight two largely overlooked aspects. First, most previous studies simplify patch encounters, neglecting the interaction between revisit dynamics and patch depletion dynamics; our results indicate that search outcomes depend critically on this interaction. Second, we show that interpatch distance strongly affects the number of revisits, particularly in finite spatial domains and over biologically relevant timescales. As a consequence, increased distances between patches can substantially increase cumulative residence times within individual patches, reducing the need for behavioral adjustment to patch spacing. This result also modifies a central prediction of the Marginal Value Theorem: when future revisits are possible, the optimal residence time during a given visit should be shorter than predicted by models that assume a single exploitation event (Adler and Kotar, 1999). Therefore, optimal patch-leaving decisions cannot be understood independently of future revisit opportunities.

Our results suggest that the increase in number of revisits with inter-patch distance does not arise from systematic changes in out-of-patch behavior. Indeed, when looking at worm movement outcomes after exiting food patches, we found little or no differences in out-of-patch movement patterns across inter-patch distances (Figure 4A, B). We also showed that the observed number of revisits are compatible with a random walk model that uses identical parameters across treatments (Figure 4D). We used a deliberately simple random walk to demonstrate that the observed effects do not require complex movement strategies. However, *C. elegans* locomotion is considerably more complex, exhibiting long-range directional persistence (Peliti et al., 2013), curvature modulation during runs (Iino and Yoshida, 2009; Suwazono et al., 2025), and adaptive reorientation dynamics (Salvador et al., 2014). In particular, reorientation rates are initially elevated following food removal and progressively decline during subsequent relocation phases (Gray et al., 2005; Hills et al., 2004; Salvador et al., 2014), a behavioral transition that has been interpreted as part of a broader shift from local exploration to relocation under uncertainty (Bartumeus et al., 2016) and could therefore influence revisit probability. We did not find evidence for worms modulating their exploration intensity in response to inter-patch distance (Figure 4A), but it did increase with food quality (Figure S11).

A key factor in the increase of number of revisits with inter-patch distance is the effect of limited space. Our experimental arenas were large compared to *C. elegans*, but worms often reached the edge of the experimental plates, and this limitation seems to have played an important role in increasing revisits (Figure 4D). While this may be seen as an experimental artifact, natural environments with rich food sources are also limited in size and time, and this is especially relevant for *C. elegans*, which feeds on the bacteria growing on pieces of decaying organic matter and has a boom and bust life cycle (Félix and Duveau, 2012). For this reason, our experimental data may also be used to investigate the limits of the MVT when confronting real situations.

*C. elegans* foraging is a multi-scale phenomenon, where worms perceive a variable level of food quality and respond to it by adapting local behaviors (turning, speed, food pumping, etc.), which in turn affects patch-level outcomes (time spent in patch, probability of revisiting it, time to reach the next patch). In the present paper, we have chosen to discuss mostly those patch-level outcomes, which have the advantage of having a direct parallel with classical optimality theory and with fitness outcomes such as the total amount of food extracted from each patch. However, remaining at the patch level has two drawbacks. The first one is a loss of nuance: For example, while we claim that food patches provide diminishing feeding rate over time, this is only true on average. Worms deplete food locally within a food patch, so their feeding rate will also increase locally when they move to yet unexplored sections of the patch. An alternative description would focus on the instantaneous experience of the worm, and could apply a recent version of the MVT adapted to this perspective, and specifically developed for simple organisms (Orjollet–Lacomme et al., 2025). The second drawback is that, without looking at food intake and local behaviors, we cannot draw the link between local sensory inputs (food intake) and patch-level outcomes (patch-leaving), nor make quantitative estimations of foraging success, only discussing MVT predictions qualitatively. In order to draw this full picture, one last technical limitation remains. In our experiments, the initial geometry and quality of patches is known, but monitoring of the food distribution past this initial state was not possible. Future works should address this limitation, as it would allow foraging behavior with an unprecedented multi-scale integration.

In conclusion, our results show that even a simple organism such as *C. elegans* can exhibit foraging behavior consistent with the predictions of the Marginal Value Theorem, despite lacking explicit spatial awareness and operating under strong sensory and cognitive constraints. By combining long-term, high-throughput observations with controlled manipulations of environmental structure, we show that near-optimal behavior emerges from a combination of active behavioral adaptation—modulation of visit duration based on recent experience—and passive effects arising from the interaction between movement trajectories and environmental geometry. Importantly, our findings highlight that revisits, often neglected in classical formulations of the MVT, play a central role in shaping patch exploitation strategies in simple organisms. Together, these results suggest that optimal foraging can arise from relatively simple mechanisms, and that incorporating both behavioral and geometrical factors is essential to understand decision-making in realistic environments. More broadly, this work bridges the gap between optimality models and biological implementation, providing a framework to study how ecological constraints and organismal limitations jointly shape adaptive behavior.

## MATERIALS AND METHODS

### Worm preparation

*Caenorhabditis elegans* were all from the N2 strain, ordered from the *Caenorhabditis Genetics Center* (CGC). Strains were ordered in 2018, and stored at -80°C using standard practices (Stiernagle, 2006). Worms were reared at 22°C in 100 mm diameter petri dishes containing 18 mL of Nematode Growth Medium (NGM: 3 g/L NaCl, 2.5 g/L peptone, 20 g/L agar, 25 mL/L potassium phosphate buffer pH 6, 1 mM MgSO_4_, 5 mg/L cholesterol, 1 mM CaCl_2_) seeded with 800 µL of saturated *Escherichia coli* OP50 culture (see Bacterial cultures). The worms were transferred every 2-4 days, before the food was depleted. To prevent accumulation of mutations with respect to the CGC strains, we never used worms that were more than 30 generations away from our freezer stock. We achieved this by re-starting our worm population periodically, either from the frozen stock or from dauer larvae that were few generations away from the frozen stock. To minimize transgenerational effects from freezing and starvation, we always used worms that were at least 5 generations away from the frozen or dauer stock. Prior to experiments, worms were age-synchronized by bleaching and egg collection (Al-Asmar et al., 2022). Eggs were allowed to hatch in 10 mL of M9 buffer (Stiernagle, 2006) under continuous agitation (orbital shaker, Heidolph Rotamax 120, 250 rpm) at 22 °C. After 19–33 h, synchronized L1 larvae were recovered by washing and transferred to rearing plates. Behavioral assays were conducted following 48 h of growth on the rearing plate (range: 46–49.5 h).

### Bacterial cultures

*Escherichia coli* (OP50 strain) were streaked on NGM medium from a -80°C stock in 50% glycerol, incubated for a few days, and then stored at 4°C and restreaked to a new plate every 2 weeks to ensure survival. To prepare liquid cultures, we first inoculated 2-3 colonies from the fridge stocks in 5 mL of LB medium in a closed 50 mL Falcon tube, and let it incubate for 24 h at 22°C with shaking (orbital shaker, Heidolph Rotamax 120, 300 rpm). Then, we extracted 1 µL of this culture using a sterile loop, and used it to inoculate 5 or 10 mL of fresh LB, that we let incubate for 24 h in the same conditions, before using it to prepare experimental plates or breeding plates.

### Experimental plates

All of the experiments were run in 55-mm petri dishes containing 8 mL of Foraging Medium (3 g/L NaCl, 20 g/L agar, 25 mL/L potassium phosphate buffer pH 6, 1 mM MgSO_4_, 5 mg/L cholesterol, 1 mM CaCl_2_, 10 mg/L chloramphenicol and 100 mg/L novobiocin) (Escudero et al., 2023). The plates were poured while each plate laid flat on a leveled table, and stored at 22°C for 2 to 9 days before the experiments. Note that these plates do not contain nutrients for the bacteria, and contain antibiotics at bacteriostatic concentrations (determined by measuring the minimum inhibitory concentration, and checking viability after 24 h of exposure (Madirolas et al., 2023)), so that bacterial quality remains constant from the moment the bacteria are placed on the plates to the moment the experiment begins.

The day before the experiments, saturated liquid cultures of *E. coli* OP50 were washed three times with 5 mL of Foraging Buffer (3 g/L NaCl, 25 mL/L potassium phosphate buffer pH 6, 1 mM MgSO_4_, 1 mM CaCl_2_, 10 mg/L chloramphenicol and 100 mg/L novobiocin) by centrifugation for 5 min in a benchtop centrifuge (MyFuge 5, Benchmark Scientific, 5500 rpm). After the last wash, the bacteria were resuspended in Foraging Buffer, and batches with the OD_600_ required for the experiments were prepared using a spectrophotometer (Jenway 7200, Cole-Parmer, Staffordshire, UK). OD was measured directly for values above 0.1, and serial dilutions were performed to obtain lower optical densities. Bacterial concentration was double-checked using a colony-forming unit counting protocol on the same day, obtaining values that were consistent for every experimental day (Figure S13), as well as with our previous studies (Madirolas et al., 2023). A pipetting robot (OT-2, Opentrons, Long Island City, NY, USA, with custom modifications to handle agar plates) was used to place 0.75 µL drops of bacterial culture on the experimental plates. These drops were then left to dry at 22°C until the experiments were performed the following day.

### Foraging experiment

The experiments were run in foraging plates containing 3, 7, 24 or 52 0.75 µL drops of bacteria, arranged following a regular triangular grid (Figure 2B). The theoretical center-to-center distance between food patches was 4.5, 9, 18 and 27.7 mm for each condition respectively, and actual distances approximated these theoretical values with good accuracy (Figure S1I). Controls (“OD = 0”) were prepared using drops of buffer, placed in the same position as the 24-patch condition. There were 40 to 101 plates (21 to 77 post data curation) for each condition. Replicates from all conditions were recorded every experimental day, in a randomized order and location, except for some conditions that were only ran in the second half of the experimental period (Table S2). After 48 h on food, the synchronized worms were washed 6 times with M9 buffer + 0.1% Triton X100 (triton prevents them from sticking to the pipette tips), performing 5 s spins on a tabletop centrifuge (mySPIN 6, Thermo Fisher Scientific, 6000 rpm). Worms were then left in a sterile 3.5 mm diameter petri dish with 2-3 mL of M9 + 0.1% Triton X100 for 65 to 90 min. This period was added to reduce the relative difference in the time spent off food between the first and the last worms placed on the experimental plates, since this time may change their motivation for food-seeking. Individual worms were then fished with a micropipette and a 1 µL droplet containing a single worm was placed at the center of each experimental plate, outside any food patches. The plate was then covered with its lid. Each experimental round consisted of 48 plates, and 16 to 55 min elapsed between the first worm being placed on the experimental plate and the start of the recordings. The total time off food prior to the start of recordings ranged from 1.5 to 2 hours, encompassing both incubation and handling steps. The 48 plates were prepared in two sets of 24 plates, and each set started recording as soon as it was ready. Therefore, the time between a worm was placed on its experimental plate and the start of the video ranged from 8 to 28 minutes. During this time some worms moved substantially, so the first recorded position of each worm was often far from its initial one, and sometimes inside a food patch (Figure S14).

At the beginning of each recording, an LED band was slowly slided above the experimental plate, to make the bacterial patch edges visible through oblique illumination (Figure S2). Then, a white LED screen covered with a privacy filter (3M, model BPNAP004 or similar) was placed to obtain uniform illumination with adequate contrast. The plates were then recorded for 8 hours at 22°C, using a custom-made setup, built using OpenBeam (MakerBeam, Utrecht, The Netherlands) with one 8-Megapixel camera (ELP-USB8MP02G-L75, Ailipu Technology, Shenzhen,China) filming each plate, and with fans to prevent temperature increase. Videos were recorded at a framerate of 1.2 frames per second, using custom software coded in Matlab (release 2023a, The Mathworks, Natick, MA, USA).

### Tracking

Worm videos were automatically tracked using custom-made code in Matlab (Release 2023a, The Mathworks, Natick, MA, USA). Centroid trajectories were then smoothed according to the following method: let *X*_0_ be the initial position of the centroid at time t = 0. Then, as long as the centroid remains within a radius of 2 pixels around *X*_0_, any movement is considered as noise.

Let *X_t_* be the first position of the centroid that is outside of this 2 pixel radius, which happens at time *t*. Then, all the centroid positions between time 0 and time *t* are interpolated linearly between *X*_0_ and *X_t_*, and the process restarts with *X_t_* as the initial position.

### Alignment and scaling of the environment

The food patches were placed by a pipetting robot to maximize repeatability, but some variability is inevitable, so to increase the accuracy we programmed the robot to make four holes at the corners of a 32-mm side square with a clean pipette tip after placing the food patches (Figure S1A).

These four holes allow us to correct for plate rotation, and give us a coordinate reference that is better aligned to the actual position of the robot’s pipette, and therefore better aligned to the food patches. The distance between those reference points was used for pixel-to-mm conversion. There was a 2-3% variation between the plates (Figure S1F), that we deemed low enough to use the average ratio (1 pixel = 0.0324 mm) for the whole analysis.

### Detection of the food patches

Despite the high accuracy and repeatability of the pipetting robot, food patches and reference points were still slightly displaced with respect to their theoretical positions (Figure S1D). Also, different drops may have slightly different sizes, and their shape is not perfectly circular. In order to have an accurate estimation of the exact position of the edge of each food patch, we proceeded as follows.

#### Oblique illumination composite

Food patches are not visible with the illumination used to view worms. To solve this issue, at the beginning of every video we passed a sliding linear light behind the experimental plates (Figure S2A). Contrast at the edge of this light is extremely high, showing the edges of even the less dense food patches. We then built a composite image, taking the region with highest contrast of each frame during the sliding illumination (Figure S2B). We used this image to find the exact position of the edge of each patch: First, we started with the theoretical position of the patch, and placed 8 points, equispaced along its edge. Then, we used custom software to automatically correct these 8 points, moving them towards the maximum of radial contrast in our composite image, which indicates the edge of the patch.

We described the edge of each food patch using a spline in radial coordinates: We define the center of the food patch as the average position of the 8 reference points. Then, we transform the reference points to polar coordinates with respect to this center, and we fit a 4-th order spline to their radial coordinate, ensuring continuity in every point, and also ensuring periodicity. This spline gives a perfect circle when the reference points are equidistant to the center, and approximates very well the edges of most of our food patches when they were not perfectly circular.

#### Manual correction

This procedure gave excellent results in most of the food patches, but not in all of them, either because of mistakes of our contrast-detection algorithm, or because the shape of some food patches was not well described with only 8 reference points. For this reason, we also performed a step of manual correction: We selected all food patches that were closer than 100 pixels to the trajectory of the worm (i.e. all visited food patches, with a conservative margin of 100 pixels), and we reviewed them visually, adding and/or removing reference points until the spline followed the edge of the food patch with great accuracy in every point.

This procedure provided a description of the edges of every visited food patch within 1 pixel of its true position.

### Data curation

#### Missing tracks

Our tracking algorithm did not find a worm silhouette in part of the frames of the videos. This can be due to the worm exiting the tracking arena and exploring the borders of the plate, where it is not visible (true negative), or due to the worm being lost by the tracking algorithm (false negative). In the case where the worm disappeared and reappeared in the same situation (if it disappeared outside food patches, and then reappeared outside, or if it disappeared and then reappeared in the same food patch), we infer that the worm was in this situation during the whole missing track (without interpolating positional data).

In some videos however, the worm disappeared outside and reappeared inside of a food patch (or vice versa). This was especially common in the 24-patch condition, where two of the outer patches were sometimes partially out of the tracking area (so if the worm exited the patch from its untracked side, it might then reappear outside, making visit duration unclear). When those events were short (*<* 1 min), we divided evenly the missing track (so if a worm disappeared inside a food patch for 60 seconds, and reappeared outside, we count it as having spent 30 of those seconds inside, then 30 outside). When those events were any longer (>1 min), we left them blank (Figure S15). For some analyses, events who start, or end with such a missing track were omitted, due to their actual duration being unclear (Figure S16). Videos containing more than 10 minutes of blank missing tracks were excluded.

#### Double tracks

Our tracking system detected more than one worm in some frames. These extra frames could be due to a tracking error (false positive), or to an actual second worm that was accidentally added to the petri dish (true positive). In order to exclude videos with more than one worm, or too many false positives, we excluded the trajectories containing 1% of double frames or more.

#### Condensation-related errors

After six hours of videos, visible condensation sometimes formed on the lids of our plates, making the tracking unreliable. In most cases our tracking system still identified the real worm, but its contour was sometimes unreliable, for example because it included part of a condensation drop. We still used these silhouettes, because we believe that they would make our data more noisy, but not particularly biased. However, in *<* 4% of plates, a detected silhouette was so large that the same worm overlapped with two patches simultaneously. When this happened, we excluded the whole video.

#### Bad patch detection

Plates where the detected food patches overlapped with each other were excluded.

#### Short videos

Trajectories with less than 10,000 seconds (around 3 hours) between their first and last tracked frame were excluded.

#### Standardization of video duration

Some metrics, such as number of visits per food patch, depend on the total tracked time for each worm (which varies since worms can exit the tracking area for variable periods of times). When computing these metrics, we used the 25000 first seconds of tracked events in each video, and excluded any videos which did not have enough tracked time. For metrics that did not depend heavily on total duration (for example durations of individual visits), we used the full videos, and we only excluded videos shorter than 10000 seconds.

#### Summary

See Table S1 for a detailed summary of the data curation used for each figure.

### Heatmaps in idealized landscapes

The heatmaps presented in Figure 1, 2, S4 and S5 were made by mapping all of our experimental data onto idealized landscapes with perfectly round and equidistant patches. In order to do so, the position of the pixels in the experimental plate was expressed in radial coordinates relative to the closest patch edge (so that each position was defined by the angle relative to the closest patch center, and the distance to that patch’s edge, with negative distances inside food patches). This radial position was then mapped on the idealized landscape (with perfectly aligned and circular patches), so that the new position of the worm retains the same angle and distance to the edge of the closest food patch.

For the probability of presence heatmaps (e.g. Figure 1A), we transformed the silhouettes of the worms to idealized coordinates, and for each pixel we computed the total time that a worm silhouette overlapped with it. Let *τ_s,w_* be the total time that the silhouette of the *w*-th worm overlapped with the *s*-th pixel. The probability of presence for each pixel *s* of Figure 1A shows the proportion of time spent in each pixel, calculated as

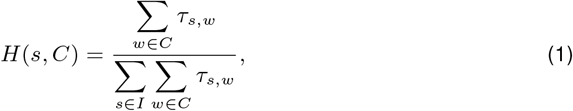

where *w ∈ C* are all the worms in condition *C* (defined by an inter-patch distance and a food quality), and *s ∈ I* are all the pixels in the idealized landscape.

For the speed heatmaps, we computed the speed of the worm centroid for each frame, and applied that speed to all pixels ovelapping with the worm silhouette. Then, for every pixel we computed the average speed for all frames in which the worm overlapped with the pixel. Finally, we averaged the speed map across individuals.

### Time spent per unit area as a function of distance to patch edge

For this analysis we again define *τ_s,w_* as the total time that the silhouette of the *w*-th worm overlapped with the *s*-th pixel. Then, we consider the distance to the closest food patch edge *d* (with negative distance for pixels inside the patch), and we define concentric rings of width Δ*d* around food patches. We call *R_d,p_* the ring containing all pixels whose distance to the edge is between *d −* Δ*d/*2 and *d* + Δ*d/*2 from patch *p*. Then, the time spent per unit area in ring *R_d_* across *visited* patches of the plate (Figure 1B, 2C and first row in Figure S6) is computed as follows

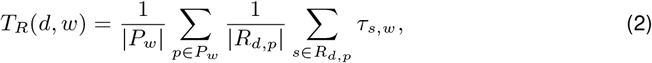

where *|R_d,p_|* is the area of ring *R_d,p_*, around patch *p*, *P_w_* is the set of all patches **visited** by worm *w*, and *|P_w_|* is the number of those patches. The previous equation provides the value for a single worm, we then averaged all the worms of each condition, and computed error bars following the bootstrap procedure described previously.

Alternative normalizations of the probability of presence

The third row of Figure S6 shows

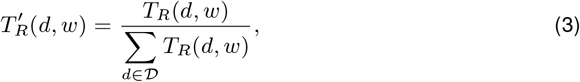

where *d ∈ D* are the centers of all the rings shown in each figure. Finally the second row in Figure S6 shows

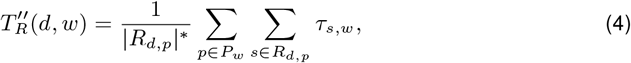

where *|R_d,p_|^∗^*is the average area of ring *R_d,p_* across all food patches of the plate. This is the same as Eq. (2) but summing food patches instead of averaging them.

### Probability and time for reaching a distance to the patch edge (Figure 4A-B)

For each worm, we extract all transits that start at the edge of a food patch, regardless of their destination (which can be the same, or another food patch, but also points where the tracking is lost, for example at the edge of the petri dish). We computed the probability to reach a given distance as *N*reach */*(*N*reach + *N*return), where *N*reach is the number of transits where the worm reaches the target distance (regardless of the final destination of the transit), and *N*return is the number of transits where the worm returns to the original patch before reaching the target distance. Note that this analysis excludes any transit that was interrupted before reaching the target distance for any other reason (i.e. interrupted tracking, reaching the edge of the plate, or reaching a different food patch). To compute the time it took to reach a given distance we took the *N*reach transits that did reach the target distance, computed the first time where the worm reached the target distance in each transit, and averaged them.

#### Random walk models

The random walk model presented in Figure 4 was implemented in R (Team, 2025), with computationally intensive parts written in C++ and integrated via the Rcpp package. Random numbers were generated in C++ using the PCG family of pseudorandom number generators (O’Neill, 2014).

We simulated individual trajectories using a two-dimensional run-and-tumble random walk. In this framework, the agent moves in discrete time steps of fixed duration Δ*t* with a constant step length *ℓ*. At each step, the movement direction (*ϕ*) is updated according to *ϕ_t_*_+Δ_*_t_* = *ϕ_t_* + Δ*ϕ*, where the turning angle Δ*ϕ* is drawn from a uniform distribution in the interval [*−π, π*]. The position is then updated as *x_t_*_+Δ_*_t_* = *x_t_* + *ℓ* cos(*ϕ_t_*_+Δ_*_t_*) and *y_t_*_+Δ_*_t_* = *y_t_* + *ℓ* sin(*ϕ_t_*_+Δ_*_t_*). Simulations were ran for a number of steps *N_s_* so that the total time (*N_s_* Δ*t*) matched the total experimental duration (8 h).

To describe the reaction of *C. elegans* to food patches, the step length depended on whether the walker was inside or outside a patch. This introduces two parameters, *ℓ*in and *ℓ*out, capturing differences in movement behavior between resource-rich and resource-poor regions.

#### Finite landscape simulations

Motion takes place within a circular arena of radius *r* = 27 mm, corresponding to the experimental Petri plates, with circular patches placed at the same positions as in the experimental landscapes and with sizes matching the measured average patch radius (*r* = 1.35 mm). Figure 4C (top) shows an example for the configuration with 7 food patches. Crossings with the edge of the arena are handled by interpolating the exact contact point with the edge and completing the remaining step with a random inward direction.

#### Infinite landscape simulations

We considered a periodic hexagonal grid mimicking the experimental arrangement but extending over an infinite domain (Figure 4C, bottom). As for the previous case, patches have the same size as in the experiments.

#### Model fitting

We fit the model parameters to each food quality in the finite landscapes. To do so, we systematically explored the parameter space defined by Δ*t*, *ℓ*in and *ℓ*out. For each combination of parameters, we run 2000 8-h simulations on each of the four configurations, and computed four observables: The number of visits per patch, the number of patches found, the average visit duration and the average transit time. Then, for each observable we quantified the agreement between simulations in finite landscapes and experimental data using a squared relative error, computed as (*x*_sim_ *− x*_exp_)^2^*/x*^2^, where *x*_sim_ is the result of the simulation and *x*_exp_ is the experimental value. Then, we added these squared errors for the four observables and for the four spatial configurations, and chose the parameter set that minimized this overall error (see optimal parameters in Table S3).

### Substitution model

The model presented in Figure 5 was implemented in Python 3.8, using libraries datatable, matplotlib (Hunter, 2007), numpy (Harris et al., 2020), opencv (Bradski, 2000), pandas (pandas development team, 2020), and scipy (Virtanen et al., 2020). It was structured as follows:

#### Visit and transit arrays

For each condition (defined by its each inter-patch distance and food quality), visits and transits made by all worms are pooled together. We first built a **visit duration array**, which stores visit durations, pooling visits to all patches, but separating visits based on their rank (first, second, … *n^th^*), so the first element of the visit duration array is a list containing the durations of all first visits, and so on. Then, we built a **transit duration matrix**. Each element [*i, j*] of this matrix is actually a list of all transit durations from patch *P_i_* to patch *P_j_*. Similarly, a **first transit array** is built, containing for each worm the first patch it has reached, and how long it took to reach it since the start of the recording (these are not included in the transit duration matrix, which only includes patch-to-patch transits). For the worms that found a food patch before the start of the recording, this first transit duration is set to 0. Finally, a **transit probability matrix** is built with in each cell [*i, j*] the number of transits between patches *P_i_* and *P_j_*, divided by the total number of observed outgoing transits from patch *P_i_*. Any transit that was interrupted by a blank missing track was excluded (see Data curation section).

#### Model for the simulations

The simulations are run in a markovian paradigm, where each node *P_i_* is associated to a food patch. When the agent is in a node *P_i_*, it has some probability of transiting to other nodes (including *P_i_*) given by the transit probability matrix. Once destination *P_j_* has been picked, the transition duration is randomly picked from the corresponding list in the transit duration matrix.

The agent then has some residency time in *P_j_*. If this is the *n^th^* visit to *P_j_*, the residency time is picked randomly from the *n^th^* element of the visit duration array.

#### Simulation start

A transit duration and corresponding starting food patch are randomly picked from the first transit array. This is necessary to match experimental data, as transits at the start of our experiments are typically longer than the average transit of the video.

#### Simulation duration

Simulations are run until 25000 s is reached, and any event beyond that threshold is truncated. These 25000 s match the one we use when computing these metrics on our experimental data (see S1).

#### Parameter substitution

To generate the results shown in Figure 5E, we proceed as follows.

##### Gray dots (full model)

The gray dots in Figure 5E correspond to the full model, where every condition is simulated with all its features (i.e. its own transit probability matrix, transit duration matrix, visit duration array, and first transit array). Each dot corresponds to the average of *N_sim_* = 1000 simulations, and the errorbars are computed as described below.

##### Blue bar (out-of-patch effects)

In order to generate the blue bars in Figure 5E, simulations are ran with number of food patches, transit probability matrix, transit duration matrix and first transit array from the focal condition, but with visit duration array from the baseline condition (the one with closest inter-patch distance). For example, the rightmost bar corresponds to an environment with 3 food patches, and corresponding transits between them, but visit durations are coming from the 52-patch condition. Each bar corresponds to the average of *N_sim_* = 1000 simulations.

##### Yellow bar (in-patch effects)

In order to generate the yellow bar (which is stacked on top of the blue bar) in Figure 5, simulations are ran with the visit duration array coming from the focal condition, but with number of food patches, transit probability matrix, transit duration matrix and first transit array from the baseline condition (the one with lowest inter-patch distance). For example, the rightmost value corresponds to an environment with 52 patches and the corresponding transits between them, but visit durations coming from the 3-patch condition. Each bar corresponds to the average of *N_sim_* = 1000 simulations.

##### Red bar (interaction)

The red bars in Figure 5 show the difference between the full model (gray dot) and the sum of the in-patch and out-of-patch effects. This difference is the interaction between both effects, which makes the full model to give a higher result than the sum of the two effects. When this interaction is negative, the red bar is not shown.

##### Proportion of each effect

The insets in Figure 5 show the proportion of each effect, computed as follows. Let *F*, *O*, *I* be the total time per patch computed using the Full model, Out-of-patch effects only, and In-patch effects only, respectively, computed as described above. The blue dots show *O/F*, the yellow dots show *I/F*, and the red dots show (*F − O − I*)*/F*.

### Averaging and bootstrapping

Unless specified otherwise, we performed averages by first averaging the data for each worm, and then averaging these values across all the worms in the same experimental condition, thus giving equal weight to each individual. Errorbars show 95% confidence intervals, computed via bootstrapping: if there are *N_worms_* worms of interest, a new set of *N_worms_* worms is sampled among them with replacement, and the metric computed over these new set. This is repeated 1000 times, and errorbars are the 2.5th and 97.5th centile of the obtained metrics. We always performed the bootstrap randomizations over the initial dataset, and for each randomization we ran all the steps needed to compute a given metric. For example, in the case of the substitution model and for a given condition, we sample *N_worms_* worms, compute all the arrays from the sampled worms (transition matrix, transit time matrix, etc.), run *N_sim_* = 1000 simulations using these arrays, and compute the average total time per patch of these *N_sim_* = 1000 simulations. This is one bootstrap replicate. We then perform *N_boot_* = 1000 of these replicates and use them to compute the confidence interval.

The only exception is the shaded areas in Figure 3F. In this case, bootstrap was performed by mixing all visit events from all worms. Then, these visits were binned by the duration of the transit preceding them. For each bin *i* containing *N_i_* visits, *N_i_* visits were sampled with replacement and their average computed. This procedure was repeated 1000 times, obtaining 1000 averages. Solid lines show the average of these averages, and shaded areas their 2.5th and 97.5th centile.

## CRedi T AUTHORSHIP CONTRIBUTION STATEMENT

**Alid Al-Asmar**: Conceptualization, Data curation, Formal analysis, Investigation, Methodology, Software, Validation, Visualization, Writing Original Draft, Writing Review & Editing. **Roger Lloret-Cabot**: Conceptualization, Formal analysis, Investigation, Methodology, Software, Validation, Visualization, Writing Original Draft, Writing Review & Editing. **Alfonso Pérez-Escudero**: Conceptualization, Methodology, Software, Resources, Validation, Writing Original Draft, Writing Review & Editing, Supervision, Project administration, Funding acquisition.

## DATA, MATERIALS, AND SOFTWARE AVAILABILITY

All the code for the analysis is available and commented in the following Github: https://github.com/martees/Video_analysis_PhD_2022.

## ACKNOWLEDGEMENTS

Harry F. Suter performed preliminary experiments that were essential to design this study. A.A. acknowledges support from a MESRI PhD grant through the SEVAB doctoral school (Toulouse), and a postdoctoral grant from the Fyssen Foundation, Paris. R.L.-C., acknowledges support from the Spanish Government through the predoctoral fellowship BES-2017-079643. A.P.E. acknowledges support from a CNRS Momentum grant, a Fyssen Foundation Research Grant, and grant ANR-22-CE02-0002 (ForAnInstant) from the Agence Nationale de la Recherche (ANR), which included a postdoctoral contract for R.L.-C. Some strains were ordered from the *Caenorhabditis Genetics Center* (CGC), which is funded by NIH Office of Research Infrastructure Programs (P40 OD010440). We thank Frederic Bartumeus, Jacques Gautrais, Jonathan Friedman, Simon Benhamou, François-Xavier Dechaume-Moncharmont, Greg J Stephens, Serena Ding, Audrey Dussutour, Julie Batut, and all members of the IVEP team (CRCA) for fruitful discussions and feedback.

## SUPPLEMENTARY METHODS

**Table S1:** Data curation summary. Green checkmark means that the criterion in the column was applied to the figures in the row. Red empty square means that the criterion was not applied. “Visits” are events that start when any pixel of the worm enters a patch, and end when all pixels have exited. “Transits” are events that start when the worm exits a patch, and end when any pixel of the worm enters any food patch.

| Plot name | Figure panels | Videos with less than 10 000 s between first and last tracked frame, any teleportation event, and > 1% of double frames were excluded | Invalid missing tracks < 1 min were shared 50/50 between transit and visit | Videos with > 10 min of cumulated long (> 1 min) invalid missing tracks were excluded | Looked at the first 25 000 s of events | Censored events were excluded (1st & last events too) | Patches with a censored visit were excluded |
| --- | --- | --- | --- | --- | --- | --- | --- |
| Heatmaps | 1A, 1C, 2B | ✓ | N/A | ✓ | □ | □ | □ |
| Distance to edge | 1B, 1D, 2C, 2D | ✓ | N/A | ✓ | □ | □ | □ |
| Visit vs time in patch | 1F, 2E | ✓ | ✓ | ✓ | □ | ✓ | ✓ |
| Time per patch | 2G | ✓ | ✓ | ✓ | ✓ | □ | □ |
| Nb visits per patch | 3B | ✓ | ✓ | ✓ | ✓ | □ | □ |
| Average visit duration | 3C | ✓ | ✓ | ✓ | □ | ✓ | □ |
| First visit | 3E | ✓ | ✓ | ✓ | □ | ✓ | □ |
| Visit vs previous travel | 3F | ✓ | ✓ | ✓ | □ | ✓ | □ |
| Cumulative sum of successive visits | 3D | ✓ | ✓ | ✓ | □ | ✓ | ✓ |
| Experimental data used in the models | 4, 5 | ✓ | ✓ | ✓ | ✓ | □ | □ |

Description of the columns in Table S1
First 3 columns: See Main Text Methods.
Looked at the first 25 000 seconds of events: About why cutting the videos, see Main Text Methods. Tracked time steps were analyzed until their cumulated duration reached 25 000 seconds. Worms without enough time steps were excluded.
Censored events were excluded (1st and last events too). Any event (visit or transit) that started or ended during a long missing track (> 1 min) or at the start / end of the video was excluded from the analysis. This is to avoid bias in the duration of events (since censored events are expected to be shorter, or even cut in half, by missing tracks).
Patches with a censored visit were excluded. For patch-level analysis, the presence of any missing track inside the patch means that we missed time that was spent in this patch. To avoid the associated bias, we simply excluded such patches from some analyses.
Description of the rows in Table S1
Heatmaps: heatmaps were shown for the full videos, as they were meant for data visualization and not for any specific analysis. Invalid missing tracks are not relevant, as they do not include the worm silhouettes used to generate the heatmaps.
Distance to edge: This is a time-step based analysis, so censorship is not relevant. Invalid missing tracks are not relevant, as they do not contain position data. Visit duration vs time in patch: We look at the full visit dynamics for food patches, so in order to avoid biases, we exclude any patch with censored events.
Description of the columns

**Table S2:** Number of replicates for each condition, after data curation. The experiments were made in two 1-month sessions, separated by two weeks, in October-November 2022. The distance are named from closest food patches (“close”) to furthest ones (“superfar”). Some conditions were only added for the second session. *: The same trajectories were used for all the controls, adding virtual food patches for each inter-patch distance. Given that the data curation depends on the position of the food patches, variable numbers of control plates were excluded for each inter-patch distance.

| Bacterial density | Distance | Number of replicates | Experimental dates |
| --- | --- | --- | --- |
| Control | close | 71* | Session 1 + Session 2 |
|  | med | 21* | Session 1 + Session 2 |
|  | far | 72* | Session 1 + Session 2 |
|  | superfar | 75* | Session 1 + Session 2 |
| OD = 0.2 | close | 39 | Session 1 + Session 2 |
|  | med | 59 | Session 1 + Session 2 |
|  | far | 77 | Session 1 + Session 2 |
|  | superfar | 36 | Session 2 |
| OD = 0.5 | close | 59 | Session 1 + Session 2 |
|  | med | 57 | Session 1 + Session 2 |
|  | far | 81 | Session 1 + Session 2 |
|  | superfar | 39 | Session 2 |
| OD = 1.25 | close | 29 | Session 2 |
|  | med | 76 | Session 1 + Session 2 |
|  | far | 39 | Session 2 |
|  | superfar | 37 | Session 2 |

**Table S3:**
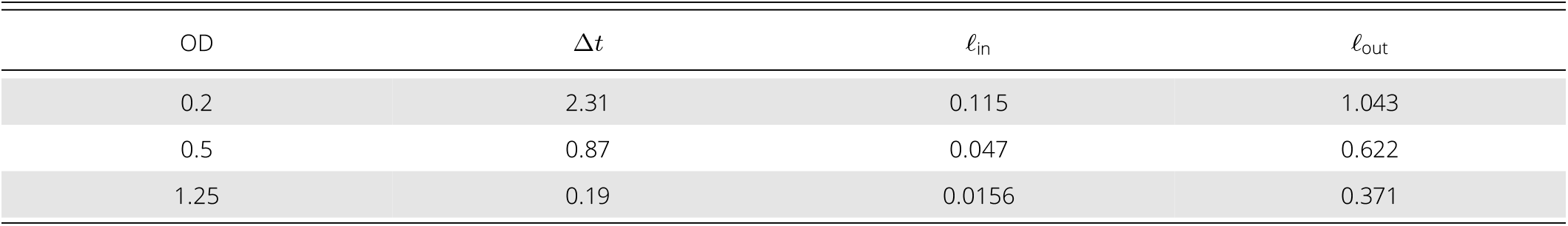
Optimal parameter values of the random walk model presented in Figure 4, obtained by minimizing the least-squares error between simulations and experimental observations for the random walk model. Δ*t* is expressed in seconds, and both *ℓ*_in_ and *ℓ*_out_ in mm.

**Figure S1:**
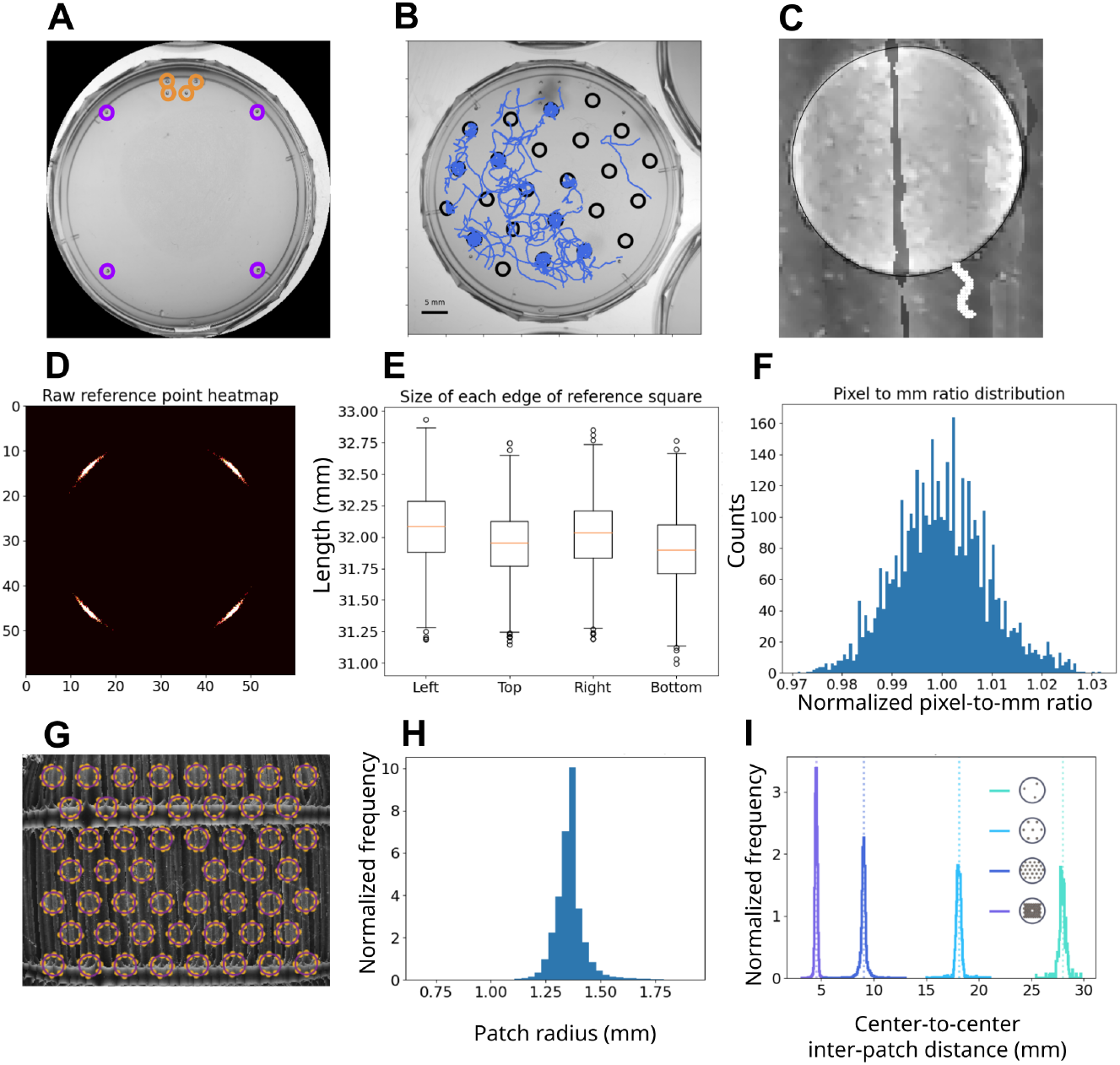
**A:** Image of an experimental plate, highlighting the points left by the pipette of our pipetor robot. The orange circles are around the binary code that allows us to automatically classify plates according to condition. The purple circles are around the four reference points that are used for position adjustment inside the plate (rotation, and scaling), to correct small errors in robot position. **B:** Example of plate with the traced food patches in black and the worm trajectory in blue. **C:** Example of tracked food patch (black) and worm silhouette (white) on a closeup to a composite image from oblique illumination (Figure S2). **D:** Heatmap of the position of the reference points in the plates, before alignment. Most of the variation lies in the rotation of the plates. **E:** Box-plot of distances between the reference points in our plates for the four edges of the square that they form. **F:** Distribution of relative pixel-to-mm ratio in all the plates, computed using the reference points (1 corresponds to the average value, 1 pixel = 0.0324 mm, used for all plates in our results). **G:** Traced patch contours (purple) and their reference points (orange) on the composite image from oblique illumination (Figure S2). **H:** Distribution of patch radii. Each datapoint in the histogram is one of 100 radii taken for each food patch. Histogram is normalized to have an integral of 1. The average is 1.36 mm. **I:** Distribution of inter-patch distances (center-to-center) in our four types of conditions. Averages and standard deviations are: 4.5 *±* 0.1, 9 *±* 0.4, 18.1 *±* 0.4 and 28 *±* 0.4 mm.

**Figure S2:**
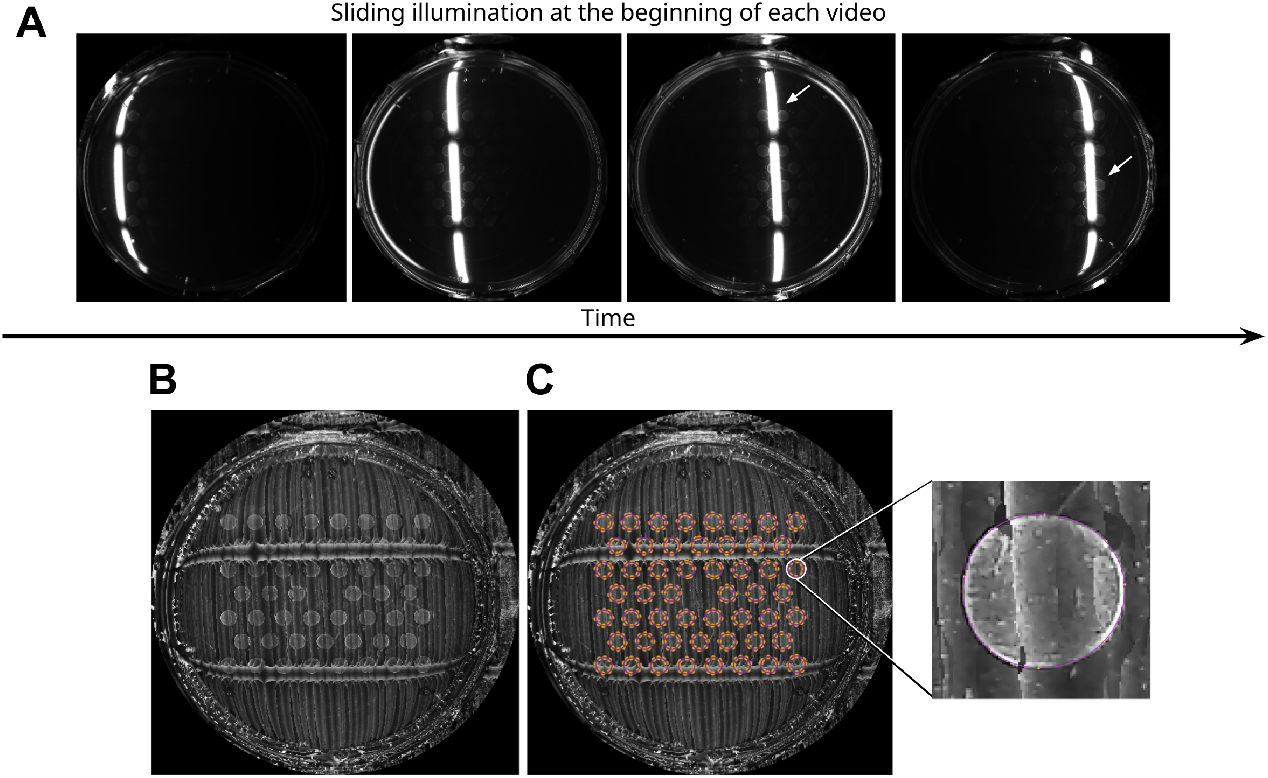
Food patch detection. In the beginning of each video, a LED band was slid behind the plates, allowing visualization of the patch edges. **A:** Four images from the beginning of the video, showing the LED band moving across one plate. The white arrows indicate places were patch edges are visible. **B:** Composite image built from the sliding illumination video. Horizontal artifacts are due to gaps in the LED band, but do not prevent patch detection. **C:** Composite image showing the splines that were fitted to the patch edges. Orange dots: Reference points from which the splines were built. Purple lines: splines fitted to patch edges. Zoom: Zoom on one patch, with the spline and orange reference points used for fitting.

**Figure S3:**
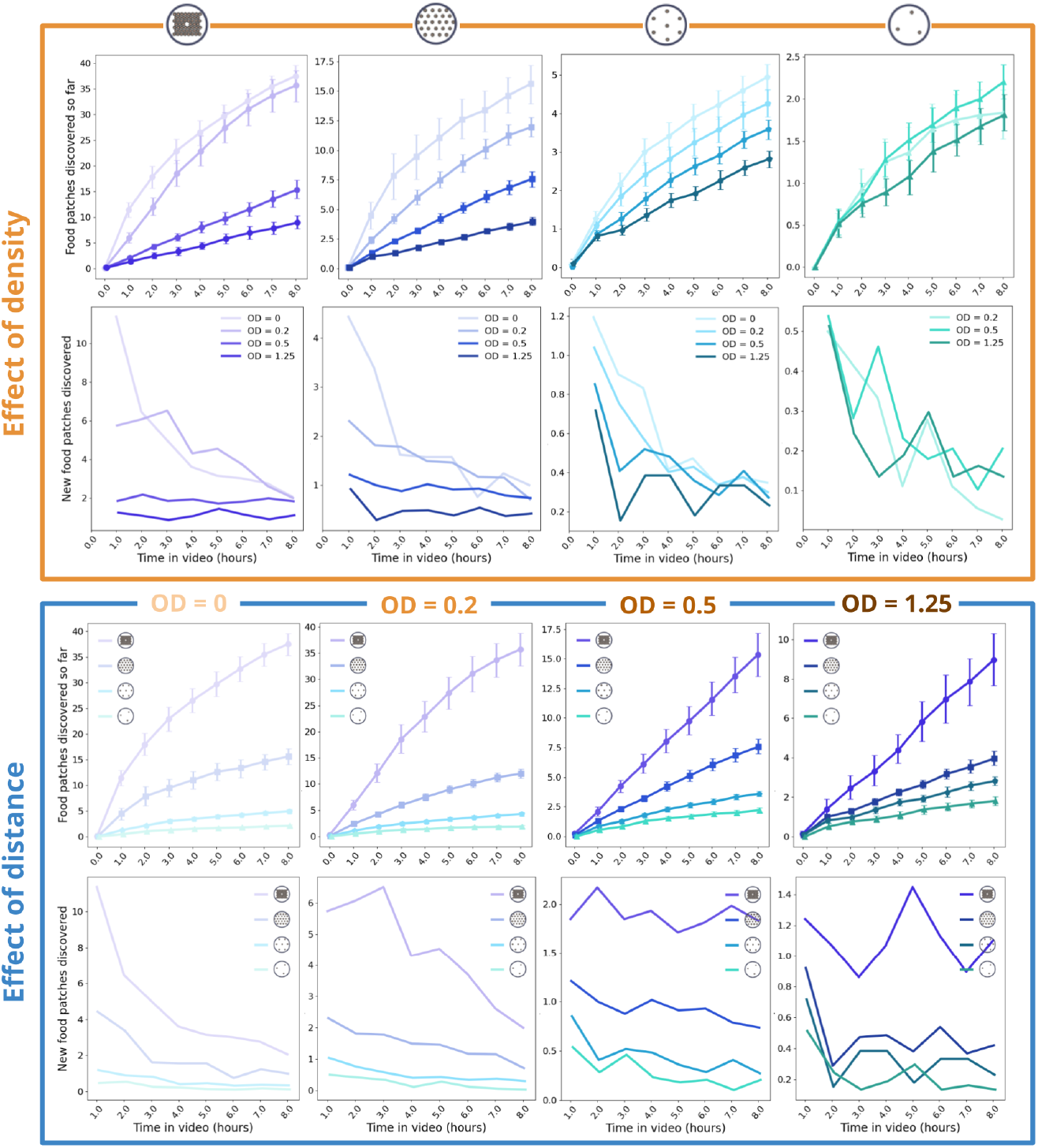
Number of discovered food patches as a function of time in the video. Orange box: curves are grouped by distance, showing the effect of quality. Blue box: curves are grouped by quality, showing the effect of distance. Each color, from green to purple, represents an inter-patch distance, and the lightness represents the quality of bacteria (lightest curves are controls, darkest curves are OD = 1.25). The first line of graphs represents the number distinct food patches discovered over time. The second line of graphs is the difference between two consecutive data points of the top line.

**Figure S4:**
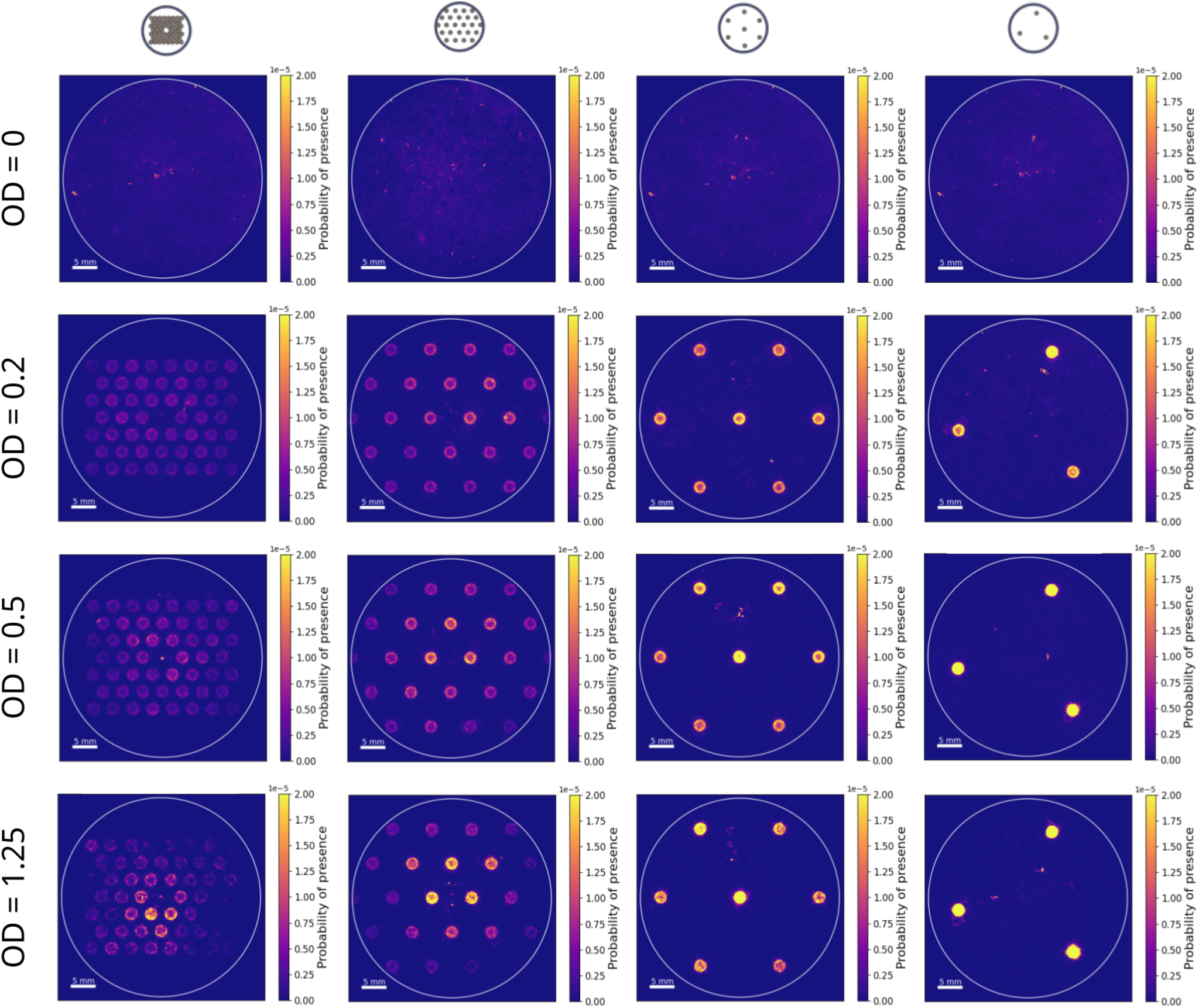
Heatmaps of probability of presence in idealized landscapes for all conditions. Each row corresponds to a food quality, and each column to an interpatch distance. These are the same heatmaps as the ones shown in Figure 1A and Figure 2B, but for all conditions. See Methods for more details.

**Figure S5:**
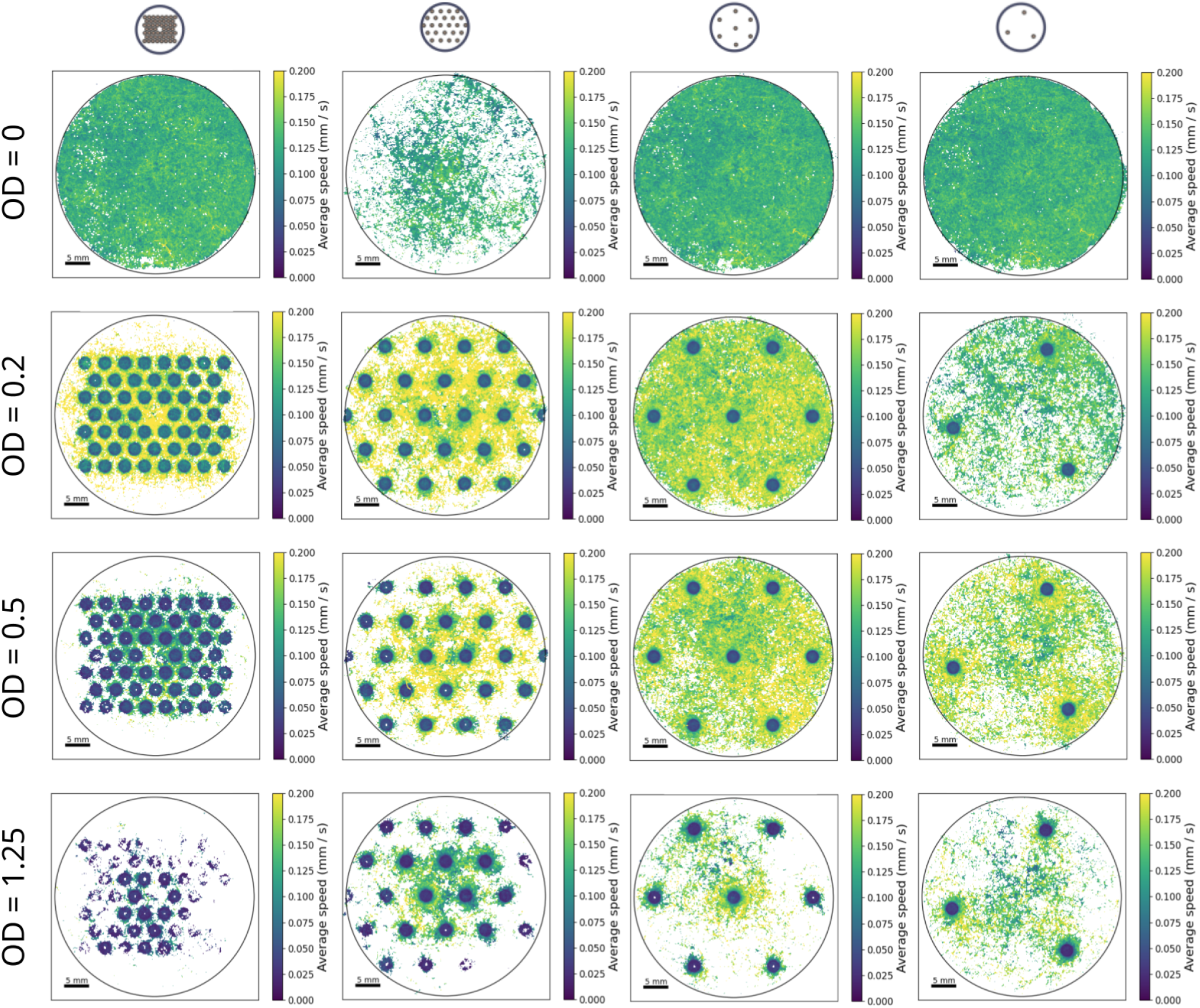
Heatmaps of speed in idealized landscapes for all conditions. Each row corresponds to a food quality, and each column to an interpatch distance. These are the same heatmaps as the one shown in Figure 1C, but for all experimental conditions. See Methods for more details.

**Figure S6:**
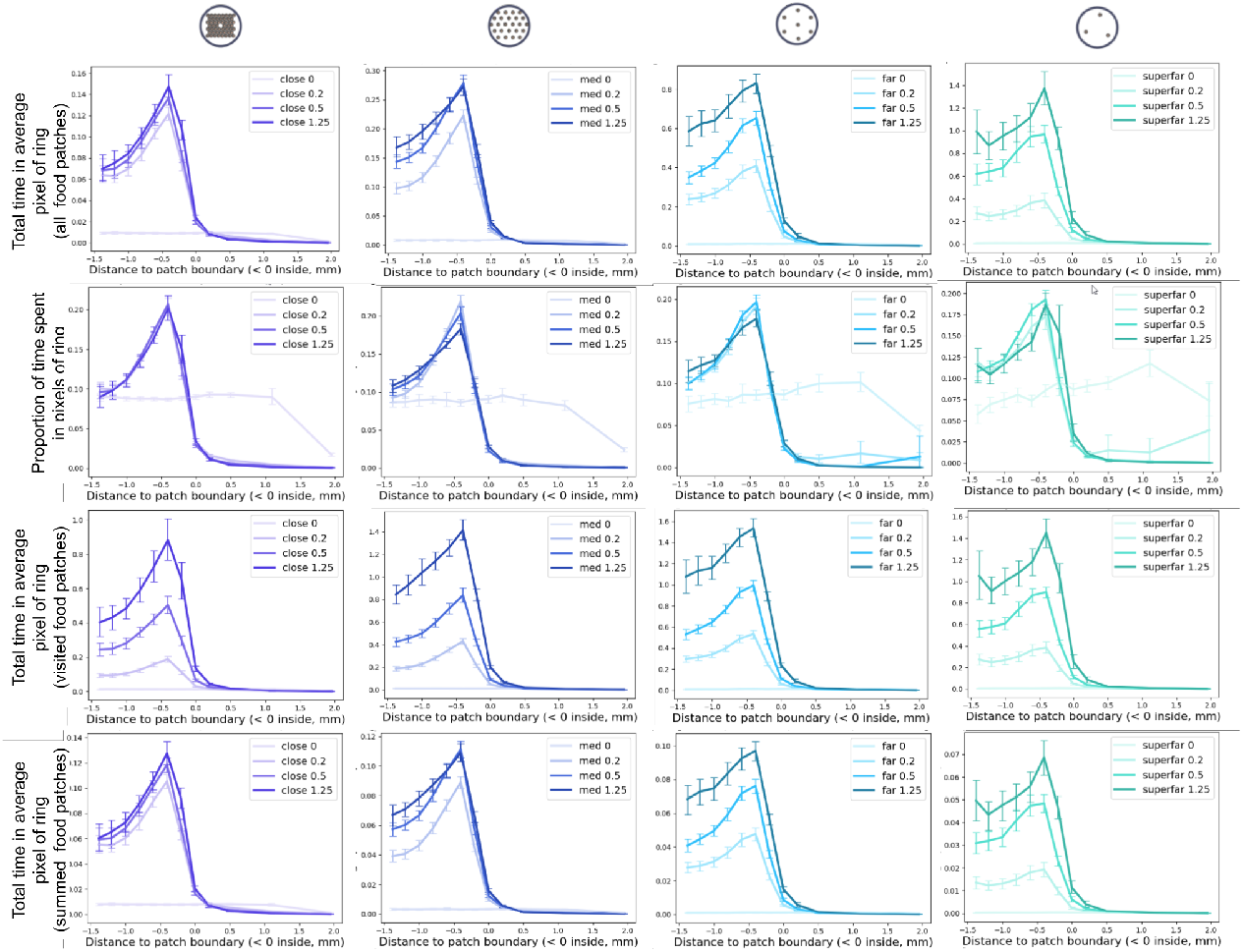
Time spent by the worm as a function of distance to patch edge. This figure shows the same results as in Figure 1B and 2C, but for all conditions, and different normalizations. Orange box: curves are grouped by distance, showcasing the effect of quality. Blue box: curves are grouped by quality, showing the effect of distance. The different normalization methods are described in the Supplementary Methods. **In both boxes: First row:** Same normalization as main text (Eq. (2)). **Second row:** Time spent per unit area at each distance from edge summed over all food patches (Eq. (4)). **Third row:** Same as first line but each curve is normalized (divided by its sum, Eq. (3)). These plots show that worms spend more time in the average food patch as quality increases (orange box, first 2 rows). They also spend more time in the average food patch as distance increases (blue box, first two rows). However, this does not reflect how much time worms have spent at that distance overall, as more patches get visited in short inter-patch distances. By summing food patches (blue box, second row), we could confirm that worms accumulate more time in patches in shorter inter-patch distances. Worms preferentially accumulate at patch border, but this tendency does not seem affected either by quality, nor by inter-patch distance (both boxes, third row).

**Figure S7:**
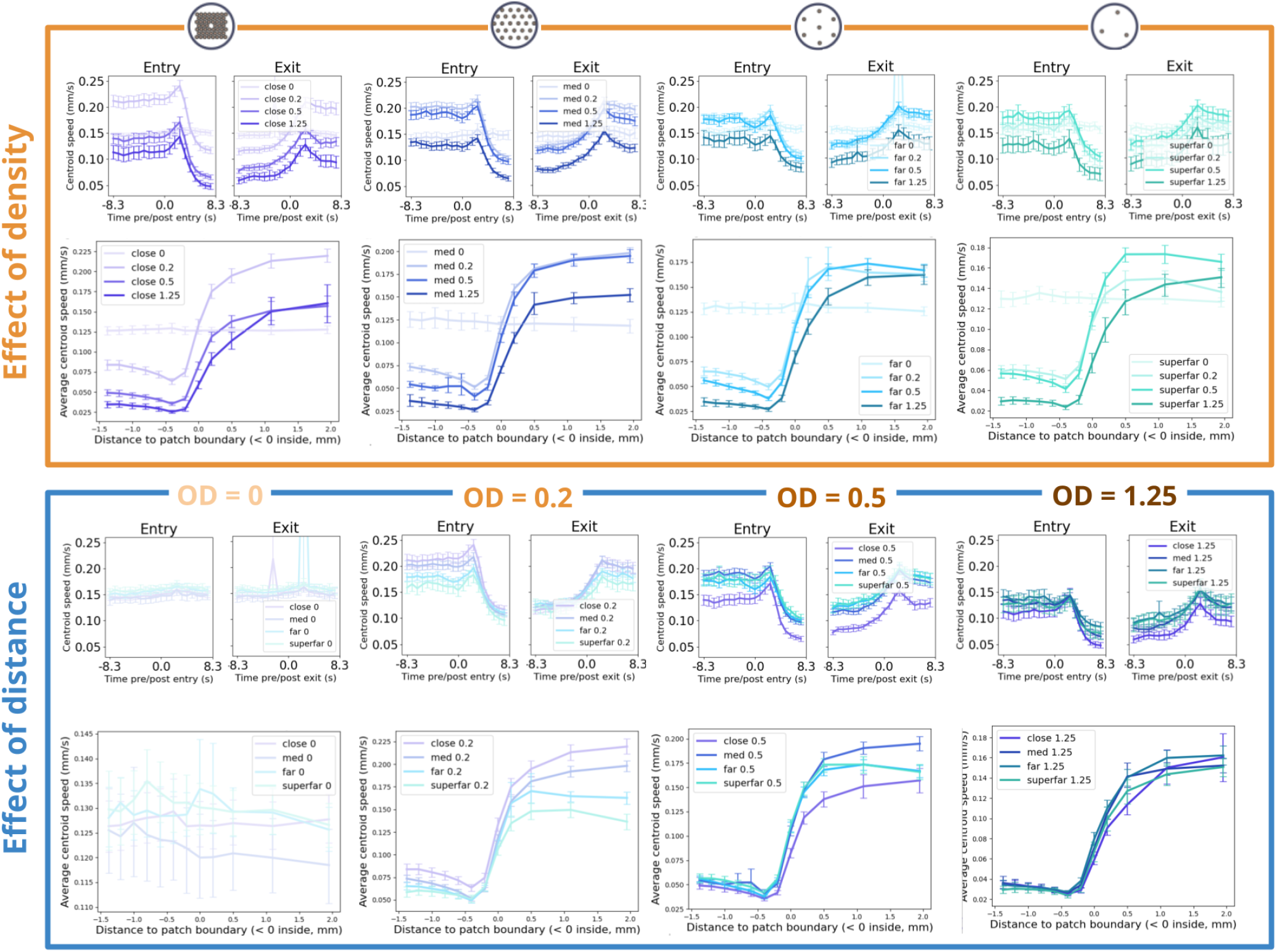
Centroid speed around patch edge in all conditions. This figure shows the same results as in Figure 1D and 2D, but for all experimental conditions.

**Figure S8:**
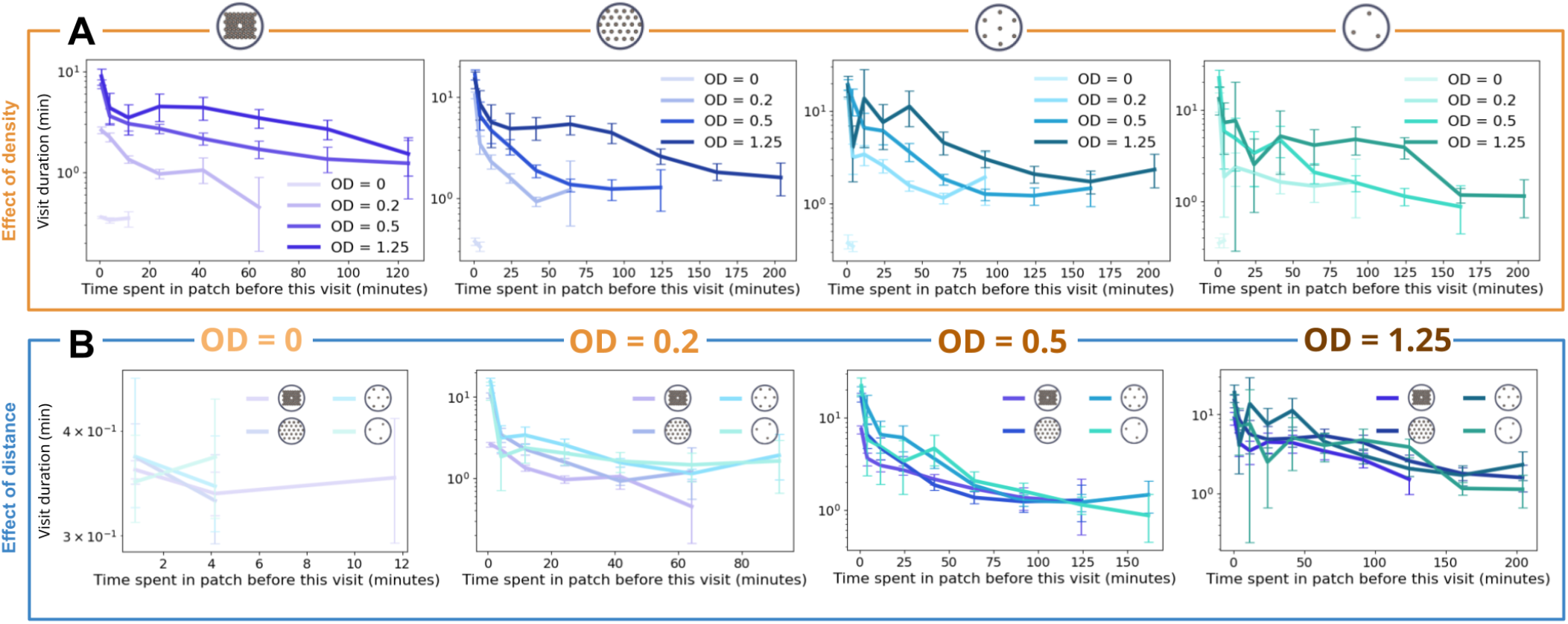
Average duration of visits as a function of time already spent in food patch, for all conditions. This figure shows the results of Figure 1F but for all conditions. **A:** Conditions are grouped by inter-patch distance, and each graph shows the different food qualities. **B:** Conditions are grouped by food quality, and each graph shows the different inter-patch distances.

**Figure S9:**
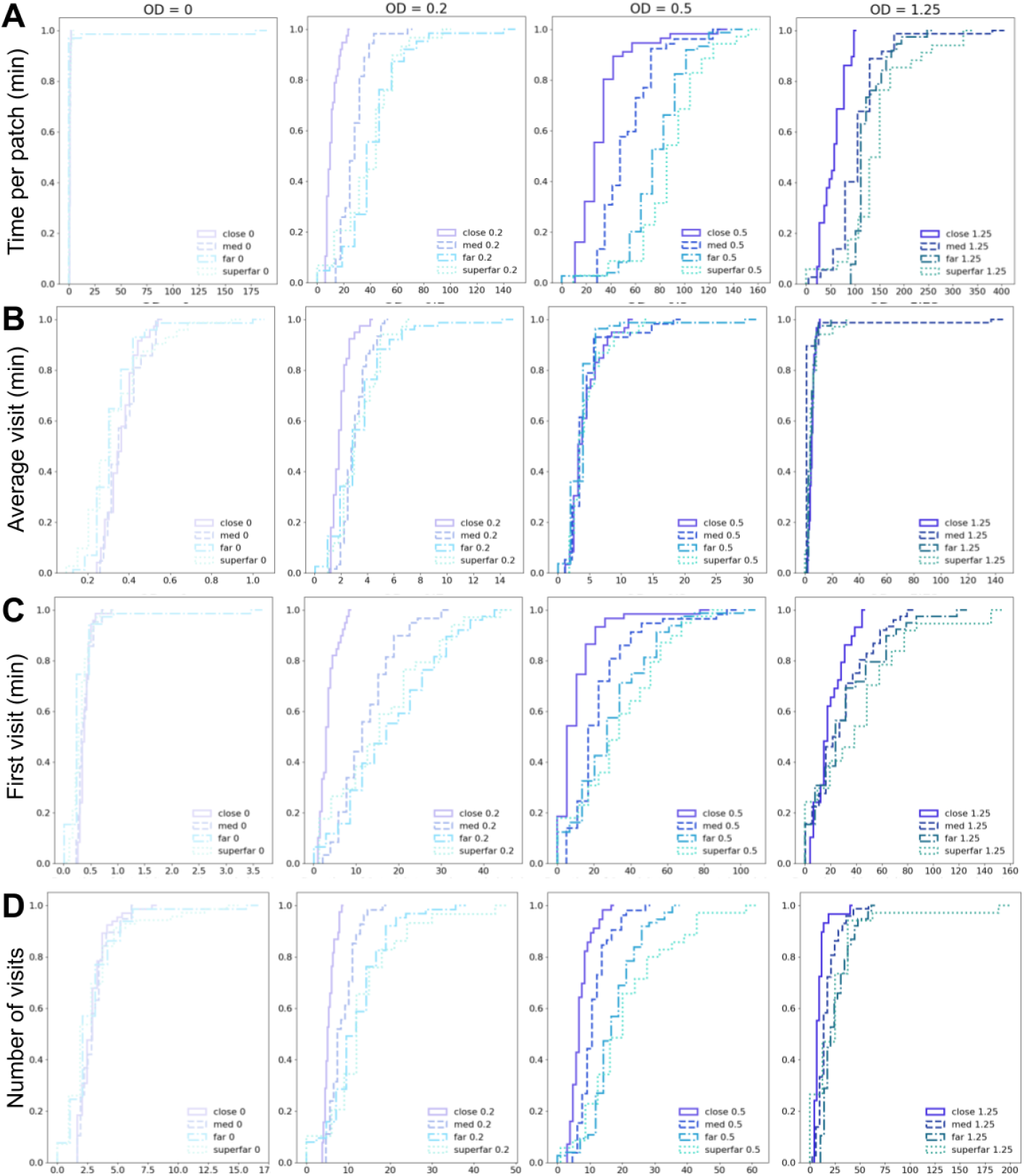
Total time, visit duration and first visit duration distribution for all conditions. The average value of each variable was computed for each worm, and then the cumulative distribution of those averages is shown. **A:** Time per patch. **B:** Time per visit. **C:** Duration of the first visit to each patch. **D:** Number of visits per patch.

**Figure S10:**
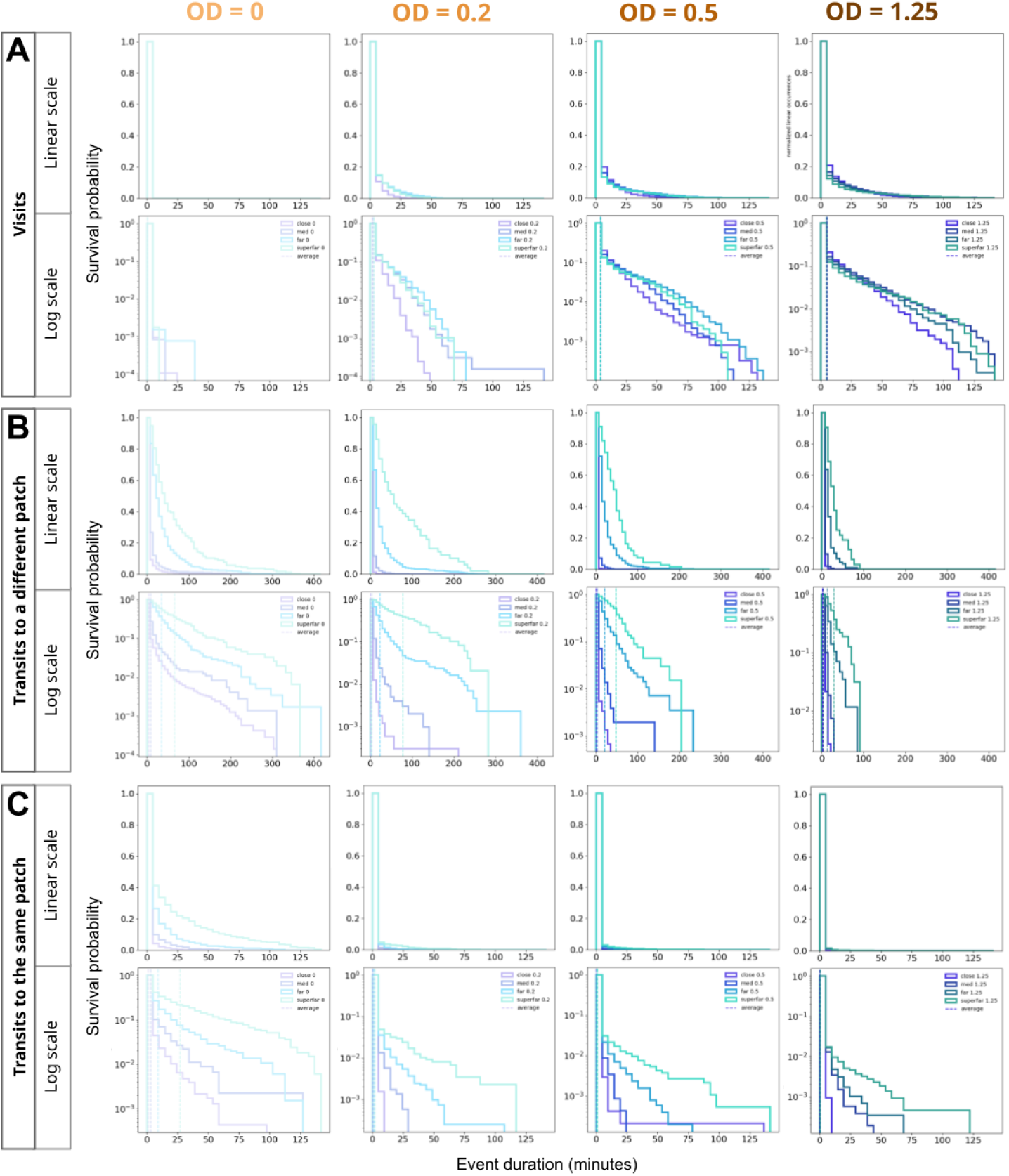
Distribution of event lengths in the different conditions. **A:** Histograms representing the distribution of visit durations, top row with a linear y-scale, and bottom row with a logarithmic y-scale. Curves represent one minus the cumulative distribution, normalized so that the first value is equal to one. This means that for each value x, the height of the histogram shows the probability that a random value picked in the distribution is higher than x. Vertical dashed lines are the average values. Conditions are grouped by bacterial quality. In each plot, one color represent one inter-patch distance. **B:** Same as A, but for transits returning to the patch the worm has just left. **C:** Same as A, but for transits going from a patch to a different one.

**Figure S11:**
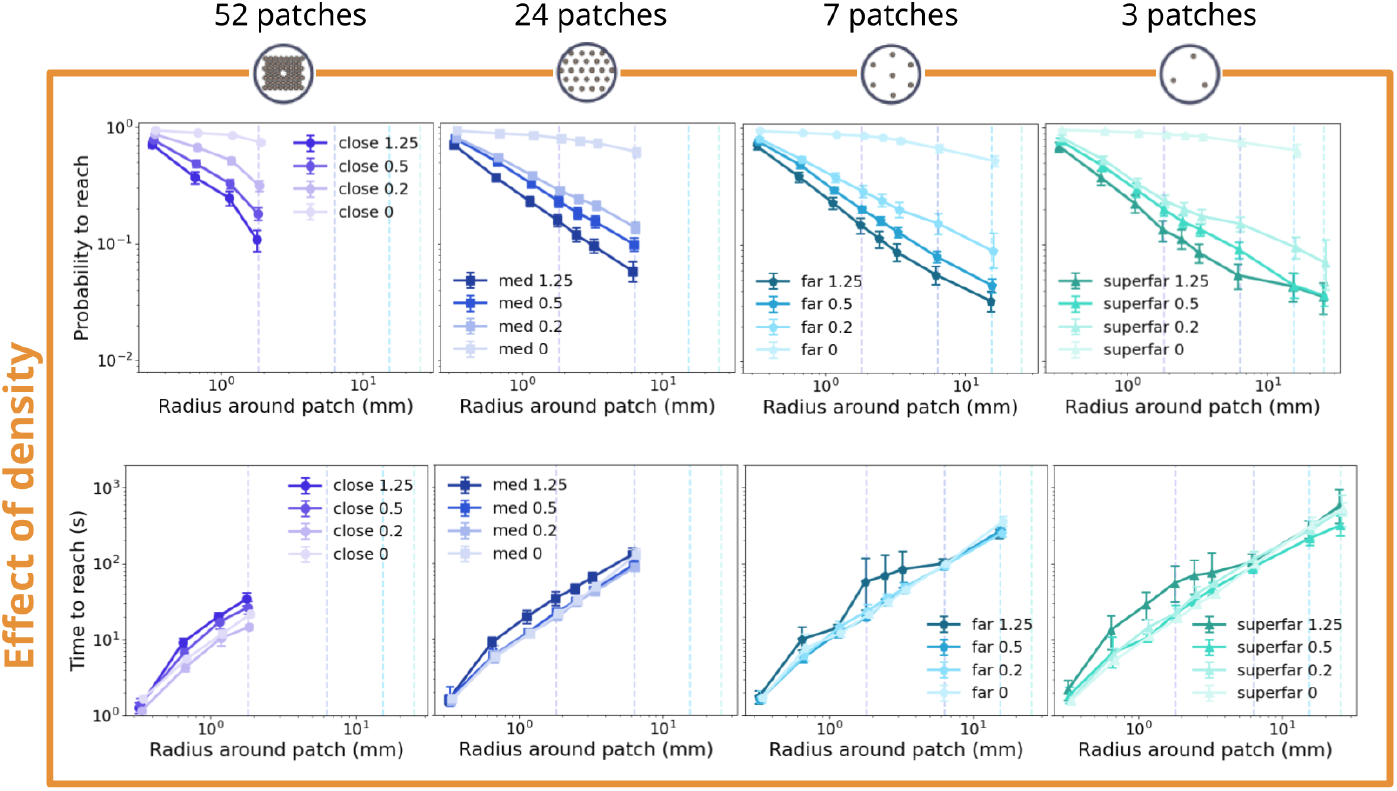
Effect of food quality on time and probability of reaching different distances from exited patch’s edge. Curves are the same as shown in 4A-B, but grouped by inter-patch distance instead of quality. These plots show that the higher the food quality, the higher the probability of worms directly returning to the food patch they have just left (as opposed to reaching further radii, and eventually a new food patch). The effect on the time to reach is less clear, but it seems that in the 52 “close” patch condition at OD = 0.5 and 1.25, as well as in the 24 “med” condition at OD = 0.5, worms do take longer times to reach each radius, suggesting that they are crawling more slowly and/or turning more often.

**Figure S12:**
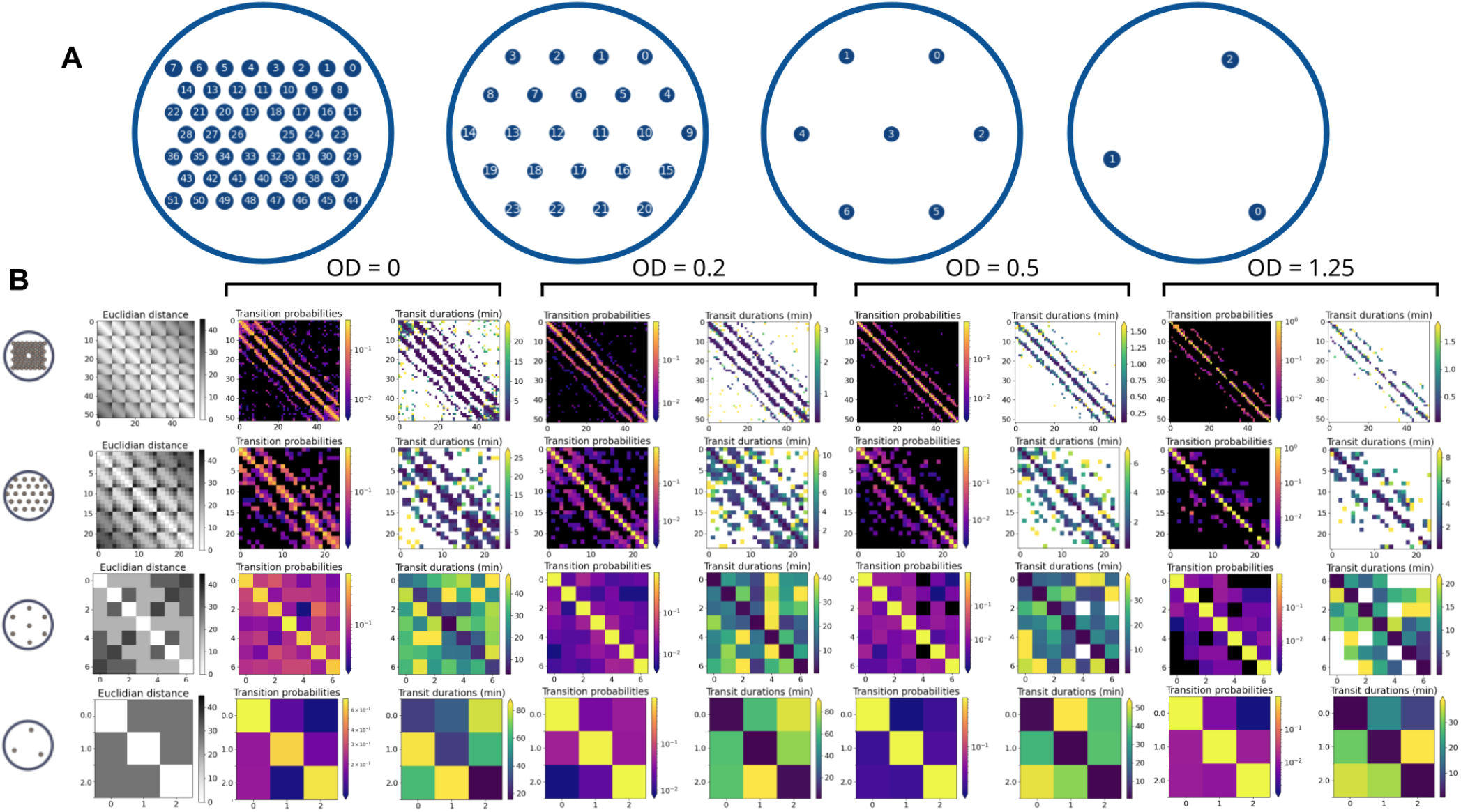
Global transit properties for our experiments. Each line of graphs corresponds to an inter-patch distance (see icons on the left). The first column represents the matrices of euclidean distances between our food patches (each cell [*i, j*] contains the distance between patch *i* and patch *j*, in mm). The second column shows the transition probability matrix for the different distances, in the controls. The third column shows the average transit time matrix for the different distances, in the controls. The other columns of graphs reproduce column 2 and 3 for the other food qualities. In OD = 1.25, some rows are white, indicating patches that were either never reached, or had no outgoing transits.

**Figure S13:**
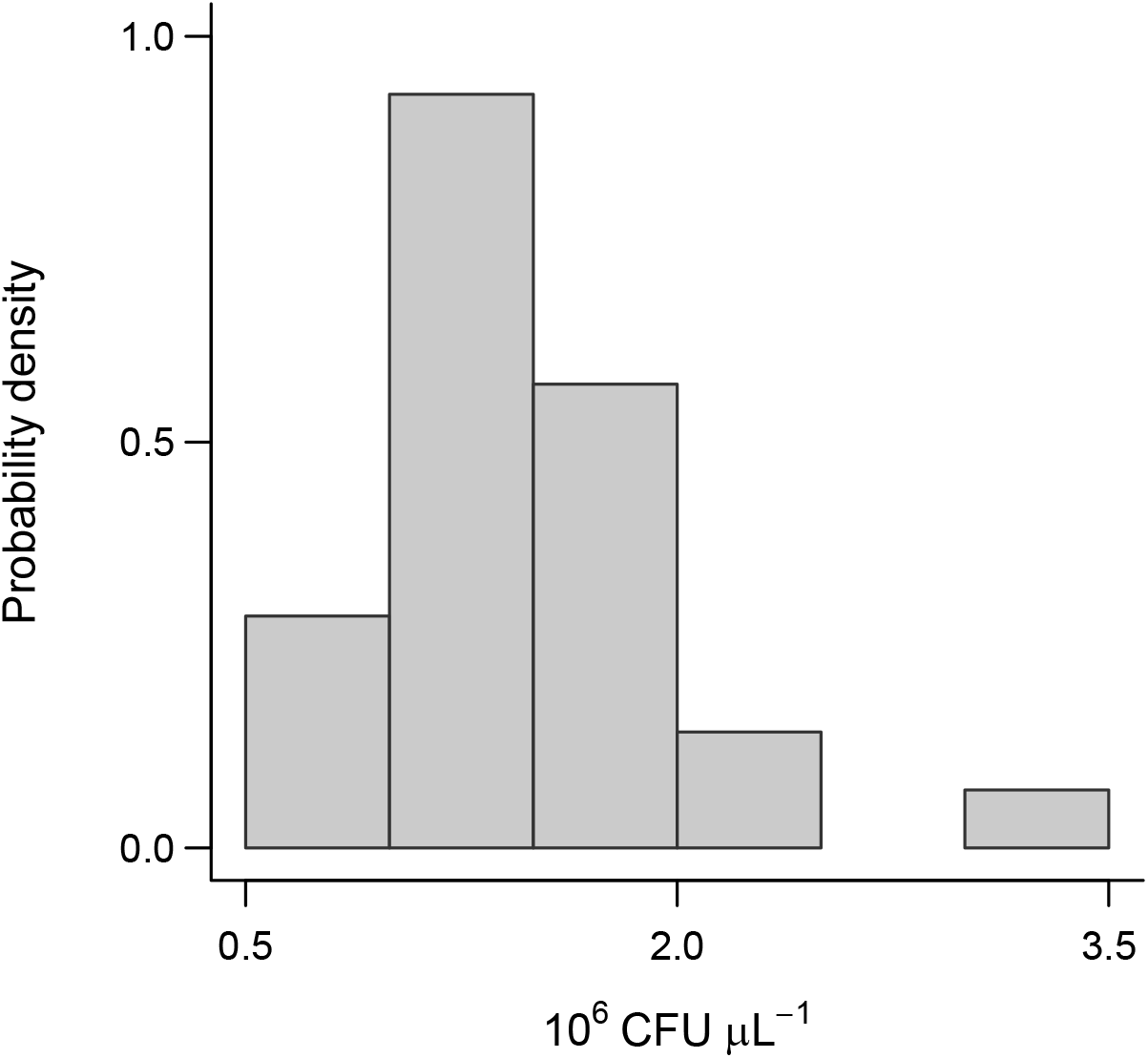
Distribution of bacterial cell density normalized to OD = 1, measured in colony-forming units per microliter (CFU/µL) across the entire experimental period. CFU counts were measured from the same bacterial cultures used to place the drops on the experimental arenas, and on the same day as the drops were placed. To measure CFU density, we performed a series of 1:10 dilutions of each bacterial culture in M9 buffer, plated four 10 µL drops from each dilution onto NGM plates, and incubated the plates overnight at 22°C. Colonies were counted from the 10*^−^*^6^ or 10*^−^*^7^ dilution, depending on colony density. CFU counts were then corrected by the measured optical density of each culture to estimate bacterial density at OD = 1.

**Figure S14:**
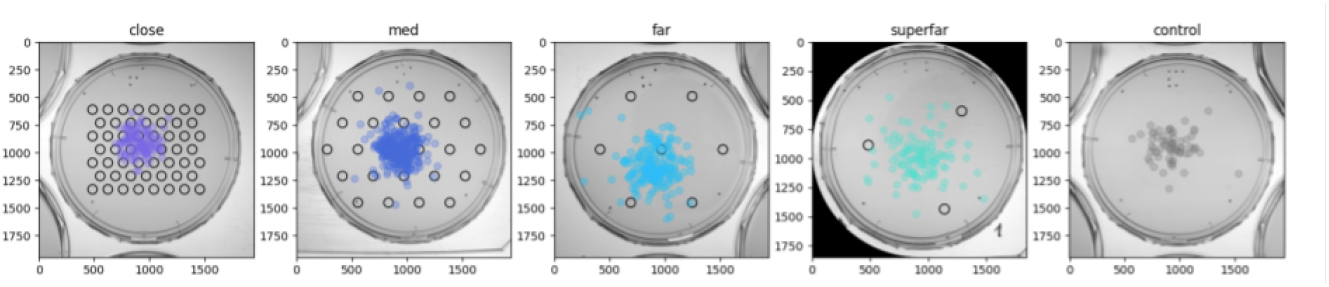
First recorded worm positions in all the inter-patch distances. All food qualities have been merged. Note that due to the position of the food patches, it was more common for individuals to start inside food patches for small inter-patch distances. Axes indicate pixels, 500 pixels = 16.2 mm.

**Figure S15:**
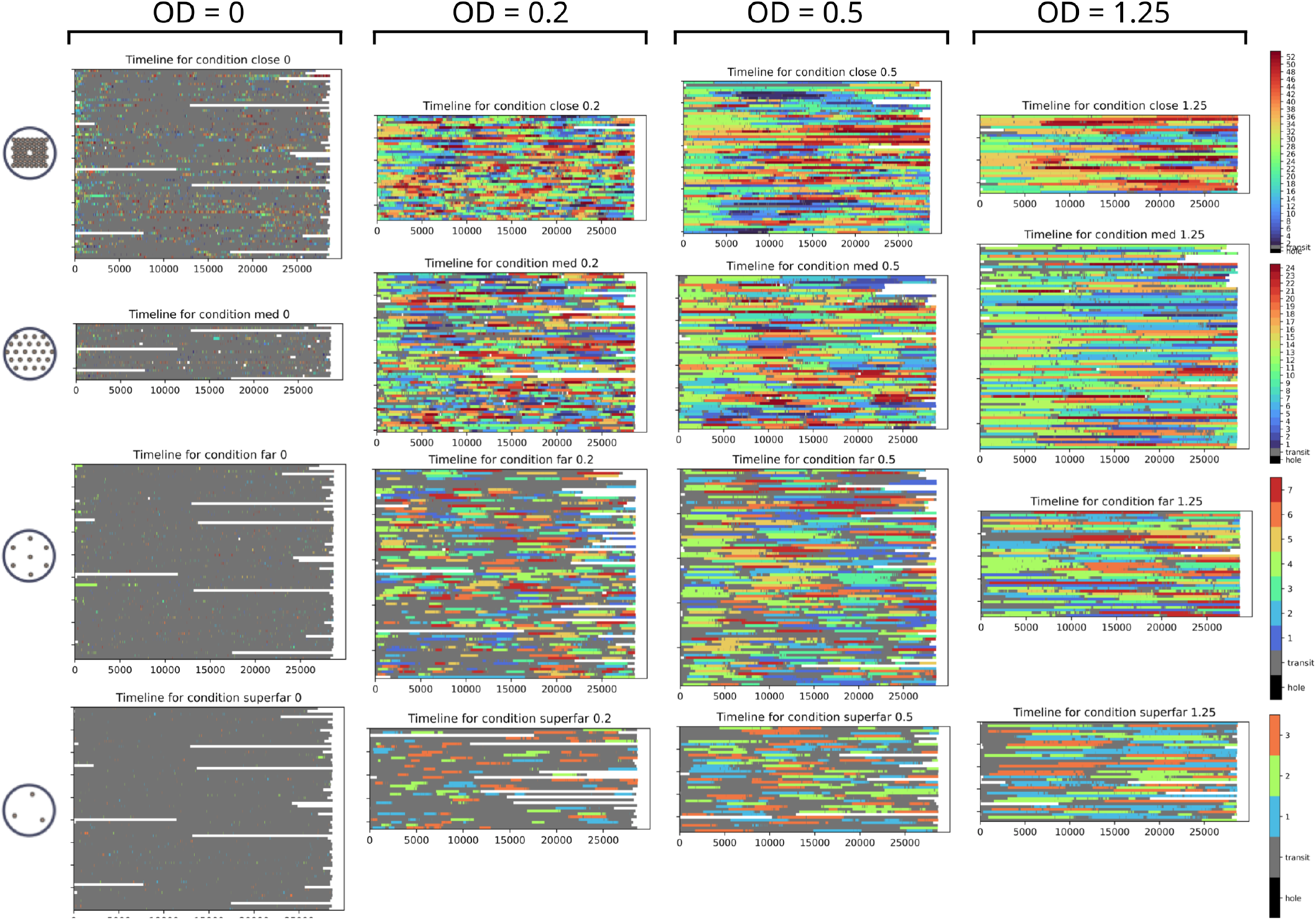
Full timeline of all of our experiments. Each row of graphs corresponds to one inter-patch distance. Each column of graphs corresponds to one food quality. In each graph, one row represents the values for one plate, and x axis is time in seconds. Color represents the state of the worm. Gray: worm is outside any food patch. White: missing invalid tracks. Other colors: worm is inside the food patch indicated by the colorbar (see numbering of food patches in Figure S12

**Figure S16:**
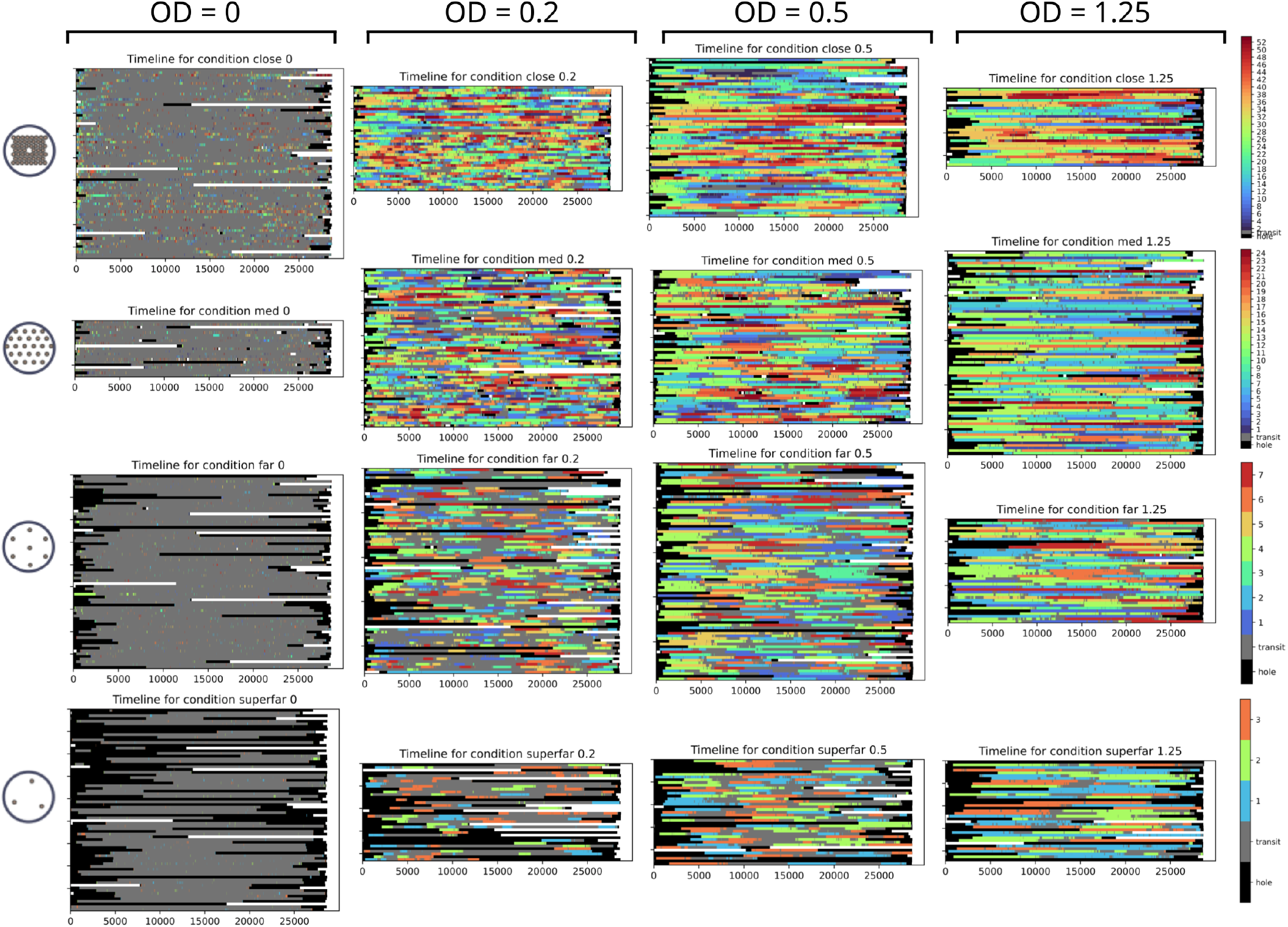
Full timeline of all of our experiments. Same as Figure S15, but showing in black the events that are considered to be “censored” during the analysis (see Table S1).

